# Exploiting public databases of genomic variation to quantify evolutionary constraint on the branch point sequence in 30 plant and animal species

**DOI:** 10.1101/2023.03.27.534366

**Authors:** Adéla Nosková, Chao Li, Xiaolong Wang, Alexander S. Leonard, Hubert Pausch, Naveen Kumar Kadri

## Abstract

The branch point sequence is a degenerate intronic heptamer required for the assembly of the spliceosome during pre-mRNA splicing. Disruption of this motif may promote alternative splicing and eventually cause phenotype variation. Despite its functional relevance, the branch point sequence is not included in most genome annotations. Here, we predict branch point sequences in 30 plant and animal species and attempt to quantify their evolutionary constraints using public variant databases. We find an implausible variant distribution in the databases from 16 of 30 examined species. Comparative analysis of variants from whole-genome sequencing shows that biased or erroneous variants that are widespread in public databases cause these irregularities. We then investigate evolutionary constraint with largely unbiased public variant databases in 14 species and find that the fourth and sixth position of the branch point sequence are more constrained than coding nucleotides. Our findings show that public variant databases should be scrutinized for possible biases before they qualify to analyze evolutionary constraint.

## Introduction

Precursor messenger RNA (pre-mRNA) splicing is executed by the spliceosome, a large ribonucleoprotein complex that assembles at the intron-exon boundary (Lee & Rio, 2015). Intronic features involved in the recognition and assembly of the spliceosome include the splice sites, polypyrimidine tract and branch point sequence (BPS). A degenerate heptamer containing the branch point residue constitutes the BPS (Keller & Noon, 1984). This motif resides within 50 bases upstream of the 3′ splice site in most of the introns across all eukaryotes. The heptamer includes highly conserved thymine and adenine residues, but the adenine itself acts as branch point during pre-mRNA splicing (Schwartz et al., 2008; Taggart et al., 2017; X. Zhang et al., 2019).

Mutations in BPS can promote alternative splicing and manifest phenotype variation (Královičová et al., 2006). However, despite their functional relevance, BPS are not readily accessible in most gene transfer files. Lack of experimentally proven branch points (Mercer et al., 2015; Pineda & Bradley, 2018) and the degenerate nature of the sequence encompassing the branch point complicate systematic annotation of this regulatory motif.

Computational methods have been developed to predict BPS (Paggi & Bejerano, 2018; Schwartz et al., 2008; Signal et al., 2018; Q. Zhang et al., 2017). Recently, Kadri et al. (Kadri et al., 2021) quantified evolutionary constraint on computationally predicted BPS in the human and bovine genomes using exhaustive variant catalogues established from whole-genome sequencing (WGS). Their analyses showed that the BPS encompasses evolutionarily conserved thymine and adenine residues that are more strongly depleted for variants than coding sequences, suggesting that they are under extreme purifying selection. Recovery of strongly constrained nucleotides within predicted BPS also shows that the motif can be localized *in silico* with high accuracy. Recent analyses in human genomes suggest that the fourth nucleotide of the heptamer might be more strongly depleted for variants than the branch point itself (Blakes et al., 2022; P. Zhang et al., 2022). However, it remains an open question if this constraint pattern is consistent across evolutionarily distant species.

Here we predict BPS in 30 plant and animal species and attempt to study their constraint using public variant databases. We uncover implausible variant distributions in 16 out of 30 public databases precluding such a study in all species. Investigation of variability of the BPS using unbiased public databases of genomic variation reveals strong evolutionary constraint on both the branch point and on the position two base pairs upstream in 14 species investigated.

## Results

Purifying selection against deleterious mutations manifests as a depletion of variation overlapping constrained nucleotides. Our previous study showed that constraints can be quantified at nucleotide resolution by counting the number of variable sites within functional classes of annotations (Kadri et al., 2021). We hypothesized that this approach is applicable to validate and quantify evolutionary constraint on the branch point sequence (BPS) for any species for which an annotated reference genome and a large and unbiased variant database are available.

### Bovine public variant database is biased

We conducted a proof-of-concept study with 89,118,442 biallelic SNPs from the bovine EVA database (release 4; (Cezard et al. 2022)) to investigate evolutionary constraint on BPS in the bovine genome. We calculated nucleotide-wise constraint - hereafter referred to as ‘variability’ - for each position of the BPS relative to the average genome-wide density of variants per 100 bp. Contrary to findings in a catalogue of variants established through WGS (Kadri et al., 2021), the branch point was the least constrained nucleotide in the heptamer when variants from the public database were used (Figure 1C). Implausible constraint patterns were also evident for other well annotated features of the genome. For instance, we found intergenic regions to be less variable than coding regions (Figure 1A), and excessive variability at the splice sites (Figure 1B). These findings suggested that the bovine EVA database contains biased or erroneous variants.

**Figure 1.**
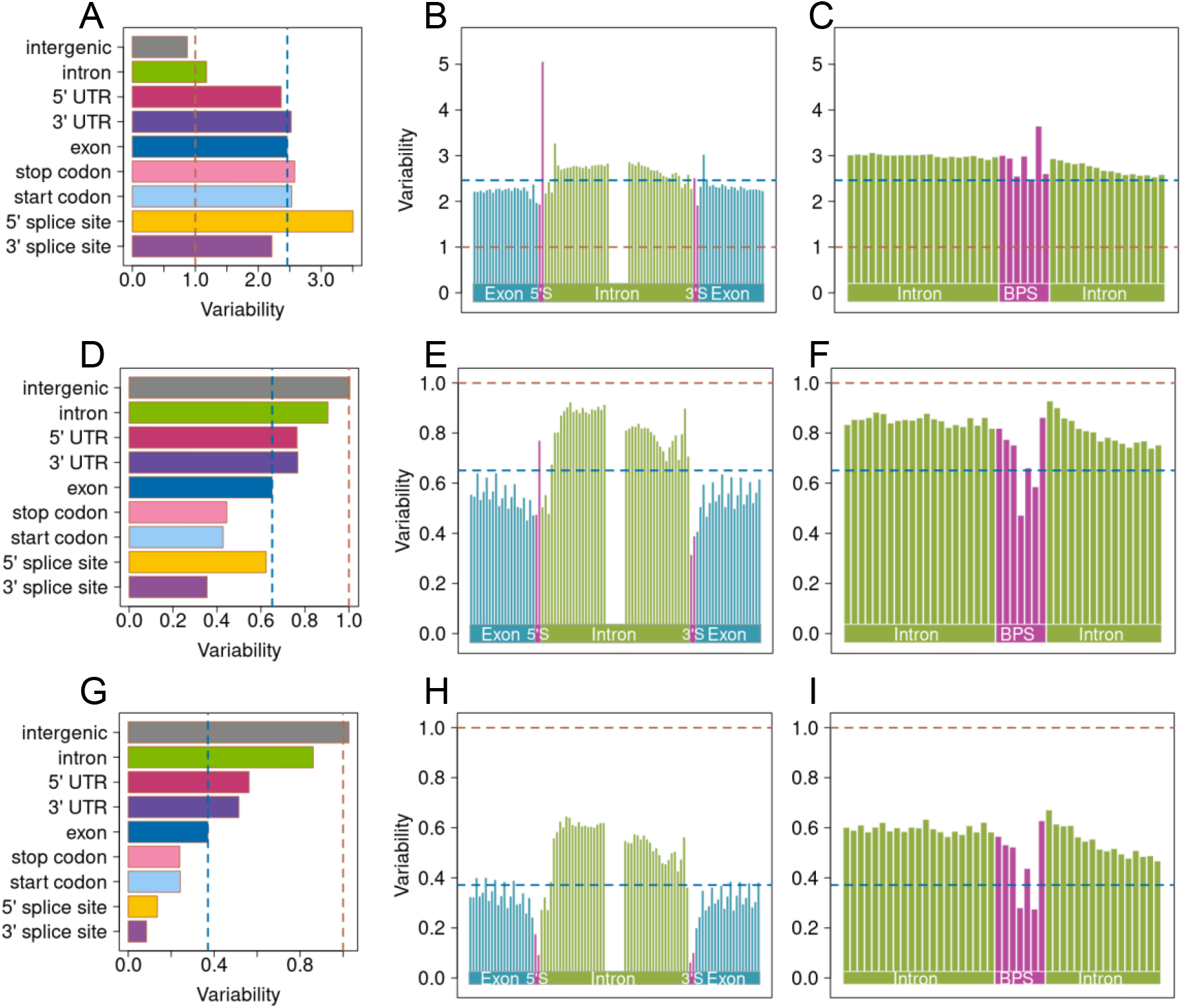
Variation in genomic features quantified through a public bovine variant database. (A, D, G) Variability of nine bovine genomic features. Nucleotide-wise constraint in and around the splice-sites (B, E, H) and predicted branch point sequences (C, F, I). Constraint was quantified relative to average genome-wide variability using (A-C) all 89,118,442 SNPs, (D-F) a subset of 57,875,698 SNPs that did not contain variants only submitted by the COFACTOR_GENOMICS_CFG20140112 project, and (G-I) a subset of 34,551,781 SNPs that contained only SNPs that were submitted at least twice. Red and blue lines denote average genome-wide and exome variability, respectively.

In fact, the proportion of coding variants was 4-fold higher in the bovine EVA database than a variant catalogue established from WGS (Table 1). An excess of coding variants in the EVA database indicated it contains variants discovered from exome sequencing. However, an implausibly low transition to transversion (Ti/Tv) ratio of 0.55 for variants overlapping exons also showed that the database is contaminated with false-positive variants. These irregularities were mainly due to a large batch of variants (n = 38,008,641) from one submitter that included many (7.45%) coding variants with very low Ti/Tv ratio (0.45). Most of these variants (83%; Supplementary Figure S1) were unvalidated, i.e., they were not confirmed by another submission. Once all variants private to this batch were removed, a subset of 57,875,698 SNPs largely recovered the expected variability of the investigated features (Figure 1DEF). However, high variability of nucleotides overlapping the 5’ splice site and a relatively low Ti/Tv ratio (1.69) suggested that this subset is still biased. We repeated the analyses with 34,551,781 variants that were submitted to EVA at least twice. Variants within this subset had a Ti/Tv ratio of 2.20. These variants recovered a pattern of evolutionary constraint that matched previous findings from WGS-derived variants (Figure 1GHI; Table 1). However, this subset contained fewer coding variants (0.49%) than the WGS-derived catalogue which suggests that strict filtration removed true rare coding variants from the data.

**Table 1.** Datasets used for analysis.

| Species | N | SNPs<br>(Ti/Tv) | % Coding<br>(Ti/Tv) | Predicted<br>branch<br>points | Average<br>density of<br>variants (per<br>100 bp) | Reference genome | Source |
| --- | --- | --- | --- | --- | --- | --- | --- |
| Cow | 266 | 29,227,950<br>(2.21) | 0.91 (3.00) | 179,476 | 1.18 | ARS-UCD1.2 | (Kadri et al., 2021) |
| Cow | - | 89,118,442<br>(1.09) | 3.27 (0.55) | 179,476 | 3.58 | ARS-UCD1.2 | EVA rs4 |
| Cow (filtered) | - | 34,551,781<br>(2.20) | 0.49 (3.33) | 179,476 | 1.39 | ARS-UCD1.2 | EVA rs4 |
| Cow (all COFACTOR) | - | 34,580,719<br>(0.55) | 7.35 (0.45) | 179,476 | 1.39 | ARS-UCD1.2 | EVA rs4 |
| Pig | 139 | 24,074,441<br>(2.36) | 0.73 (3.28) | 192,744 | 1.04 | Sscrofa11.1 | this study |
| Pig | - | 56,883,886<br>(1.96) | 0.91(2.38) | 192,744 | 2.47 | Sscrofa11.1 | Ensembl variation |
| Sheep | 161 | 16,198,123<br>(2.59) | 0.54 (4.35) | 168,025 | 0.60 | Oar_rambouillet_v1.0 | (Li et al., 2023) |
| Sheep | - | 48,541,784<br>(2.45) | 0.82(3.53) | 168,025 | 1.80 | Oar_rambouillet_v1.0 | Ensembl variation |
| Goat | 157 | 13,364,058<br>(2.48) | 0.66 (3.95) | 174,407 | 0.53 | ARS1 | (Li et al., 2022) |
| Goat | - | 31,517,363<br>(2.40) | 0.68(3.88) | 174,407 | 1.26 | ARS1 | Ensembl variation |

### Public variant databases reveal expected constraints in pig, sheep, and goat

We quantified constraint patterns in the same genomic features of three additional species to investigate if other public variant databases suffer from similar biases. These analyses were performed in pig, sheep, and goat for which exhaustive variant catalogues were available from both WGS and public databases. The predicted BPS in 192,744 pig, 168,025 sheep, and 174,407 goat introns had a “nnyTrAy” consensus sequence that contained two conserved nucleotides at the branch point itself (position 6) and two bp upstream (position 4) (Figure S2A). The branch points were at a median distance of 26 bp upstream of the 3’ splice site (Figure S2B). Most BPS had a canonical ‘TnA’ motif at position 4-6 (88, 90 and 89% in pigs, sheep, and goats, respectively).

Variant discovery in WGS data of 139 pigs, 161 sheep and 157 goats yielded 24,074,441, 16,198,123 and 13,364,058 biallelic SNPs with Ti/Tv ratios between 2.36 and 2.59 (Table 1). Variability in the genomic features differed as expected and confirmed previously established patterns of constraint (Figure 2). We observed a striking depletion of variation on the positions 4 and 6 of the predicted BPS. The constraint on both nucleotides was stronger than on coding sequences but did not differ significantly between them (Fisher’s exact test P-values 0.06, 0.79, 0.50 for pig, sheep and goat, respectively).

**Figure 2.**
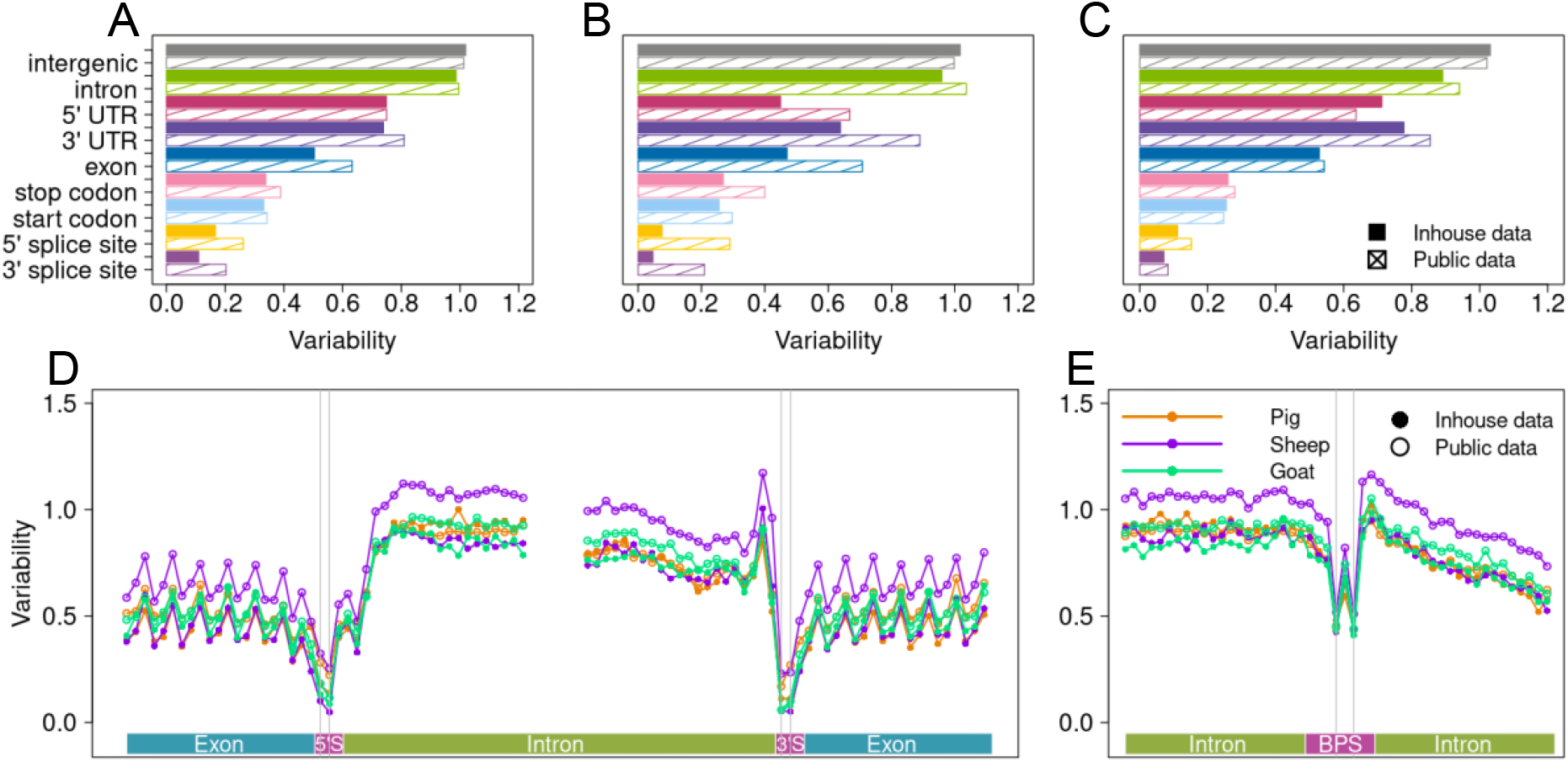
Variation in pig, sheep, and goat genomic features quantified through variants from whole-genome sequencing and public databases. Variability of the nine features of the (A) pig, (B) sheep, and (C) goat genomes. Nucleotide-wise variation relative to average genome-wide variability in and around (D) splice sites and (E) branch point sequence.

The public pig, sheep, and goat databases contained more than twice the number of variants we established through WGS but between 28% and 36% overlapped between the databases and WGS for the respective species (Table 1). Because the public databases aggregate variant information from many individuals from multiple breeds, the Ti/Tv ratios, proportion of coding variants, and constraints in functional features differed from those established with the smaller WGS subset but were within plausible ranges (Table 1; Figure 2). Nucleotide-wise constraint in the BPS was also consistent with the pattern obtained from variants established through WGS (Figure 2E). As observed with variants from WGS, the constraint did not differ between positions 4 and 6 (Fisher’s exact test P-values 0.48, 0.67, 0.47 for pig, sheep and goat, respectively).

### Variant bias is widespread in public databases

Exhaustive variant information from four public databases confirmed evolutionary constraints that were similar to those established from WGS. This encouraged us to conduct a comparative analysis of constraint on the BPS in 26 additional species (18 animals from 13 orders and 8 plants from 4 orders) for which at least one million SNPs were available through EVA (n = 8) or Ensembl (n = 18) databases. We evaluate the quality of these databases prior to the comparative constraint analyses to ensure they are free from erroneous and biased variants.

Two public databases were excluded prior to the comparative analysis because variant density was too low. Variants from 13 databases were incongruent with properties of genome-wide variants and thus were not suitable for an unbiased comparative assessment of evolutionary constraint across species (Table S1, File S1). The variability in intergenic regions was lower than the average genome-wide variability in 12 excluded databases possibly indicating biased variant distribution due to exome sequencing. An excess of exonic variants in five of these databases is further evidence that the variants fail to represent genome-wide variability. The well-established constraint at the four positions overlapping the 3’ and 5’ splice sites was absent in five databases (File S1).

Other variant characteristics such as the proportion of coding variants or the Ti/Tv ratio were not abnormal for many excluded databases, indicating that these parameters are not suitable to assess the plausibility of such databases. For instance, a Ti/Tv ratio of 1.96 and 1.11% coding variants in the *Equus caballus* database are compatible with expectations for genome-wide variants (Table S1). Yet, variants from this database revealed an implausibly high variability at both splice sites and downstream the BPS at intronic positions overlapping the polypyrimidine tract (File S1). A plausible Ti/Tv ratio (1.87) and percentage of coding variants (2.89%) may pretend that the *Gallus gallus* variant database is representative for whole-genome variability in chicken. However, an excess of variability of the nucleotides overlapping the 5’ splice sites is implausible. Variants from this database also uncovered a constraint pattern in the BPS which deviates from what we established in unbiased variant catalogues.

Only 11 public variant databases (File S1, Table S1) fulfilled our criteria, i.e., they contained on average at least one variant per 1000 base pairs, variant density was higher in intergenic regions than genome-wide (Figure 3B) and constraint on the 3’ and 5’ splice sites were evident (Figure 3A). These databases contained between 4,486,640 and 69,472,724 variants of which between 0.61% and 16.51% overlapped coding sequences. We subsequently conducted a comparative analysis of BPS variation in these species.

**Figure 3.**
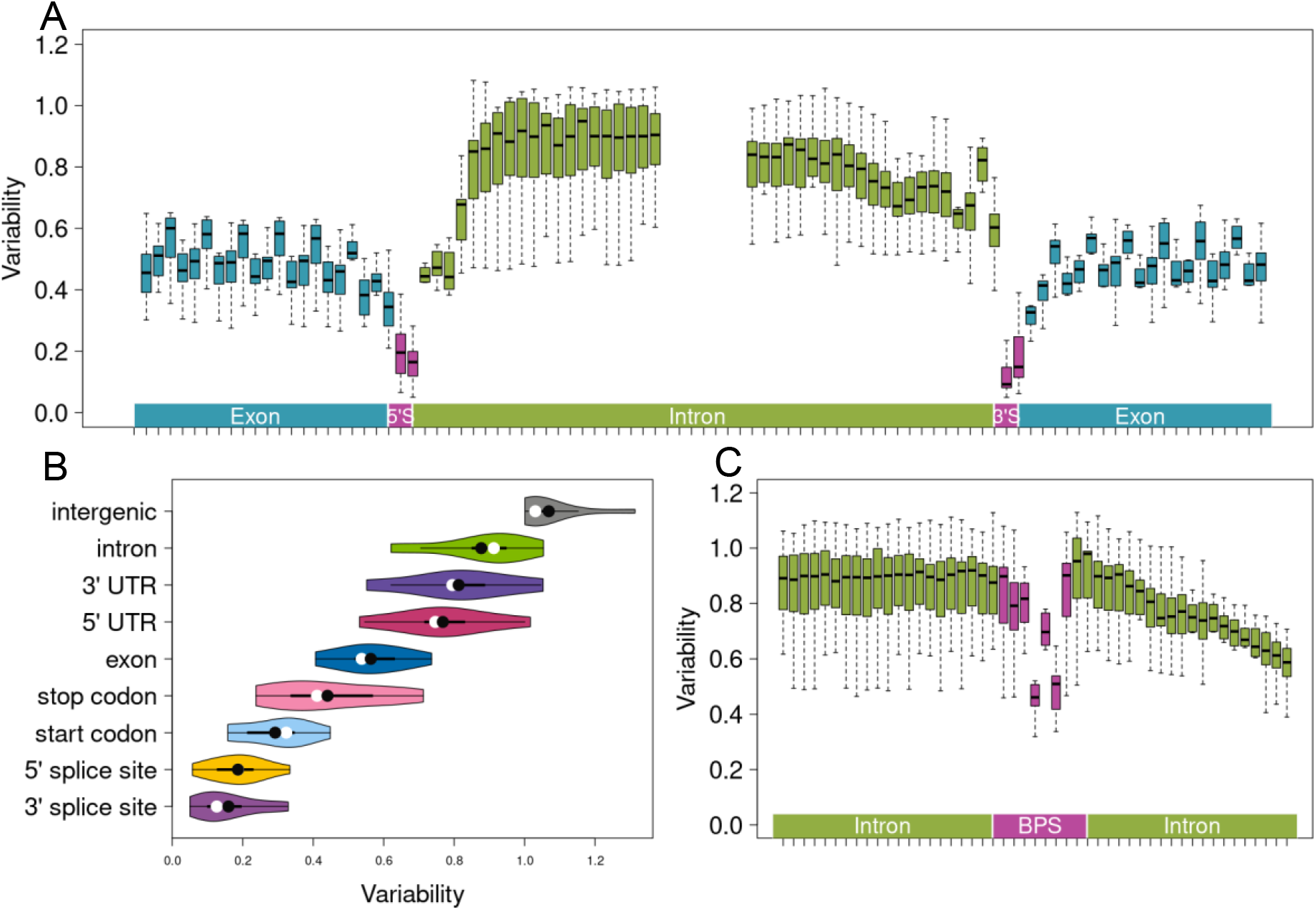
Variation in genomic features across 11 species quantified using public variant databases. Boxplots of nucleotide-wise variability relative to average genome-wide variability in and around (A) splice sites and (C) predicted branch point sequences. (B) Violin plots of variability in nine genomic features. Means and medians are indicated with black and white circles, respectively.

A vast majority of the predicted BPS for these 11 species contained the canonical ‘TnA’ motif overlapping positions 4-6 of the heptamer (between 90% in *Pan troglodytes* and 98% in *Phaseolus vulgaris*; Table S1, Figure S3). The predicted branch points were primarily between 14 and 145 bp upstream of the 3’ splice site with median distance of 27 bp (Figure S4), consistent with BPS placement in other species. The comparative analysis of BPS variation in these 11 species revealed strong constraint on positions 4 and 6 (Figure 3C, Table S1). The constraint between these two positions differed significantly in four of the 11 species investigated. In three of those four species, the constraint was stronger on the position 4 than on the position 6 (Bonferroni corrected Fisher’s exact P < 4.5 × 10^−3^).

## Discussion

Our comparative analysis of branch point sequence (BPS) variation relied on computationally predicted BPS because exhaustive catalogues of experimentally proven branch points were not available for the species considered. While the length of the reported consensus sequence encompassing the branch point varies from five bases in humans (Gao et al., 2008) to ten bases in plants (X. Zhang et al., 2019), all BPS in our study were heptamers that contained the branch point at the 6^th^ position. Constraint on positions 4 and 6 were striking in all species investigated, corroborating a pivotal role of both residues for spliceosome assembly during pre-mRNA splicing (Mercer et al., 2015). However, we did not find conclusive evidence for higher constraint on position 4 of the heptamer, as reported for human BPS (Blakes et al., 2022; P. Zhang et al., 2022). Our findings confirm that an adenine residue is the most preferred branch point, and that thymine is the most preferred residue at 2 bp upstream of the branch point (Zhuang et al., 1989). High evolutionary conservation of these nucleotides across distant eukaryotic species reiterates the need to consider them in search for trait-associated variants.

We use public variant databases to assess evolutionary constraint on different genomic features. However, contamination of these databases with erroneous or biased variants can cause flawed interpretation of results (Khafizov et al., 2015; LaDuca et al., 2017; Maffucci et al., 2019). Our study corroborates that variants from public databases need to be evaluated carefully due to their partly unknown origin and lack of curation (Johnston & Biesecker, 2013; Khafizov et al., 2015; Maffucci et al., 2019; Mitchell et al., 2004; Musumeci et al., 2010). Strict filtration, such as the removal of variants that were submitted only once, was required to recover expected constraint patterns from a public bovine variant database. This approach is only possible with accompanying metadata, which is not always available. Moreover, this approach also removes true rare variants enriched in evolutionary constraint signatures, and as such is not generally advisable.

Assessment of constraint patterns in annotated genomic features is more useful to evaluate the quality of variant databases than inspecting other widely used parameters such as Ti/Tv ratio. We show that the constraint on the splice sites and the proportion of variants in intergenic regions are the most informative for such an assessment. By using a simple and straightforward approach of counting variable sites overlapping genomic features, we show that erroneous and biased variants contaminate 16 of the 30 investigated public variant databases. Since these constraint patterns are so widespread, such an approach may provide quality assessment for existing or even purely predictive annotations.

## Material and methods

### Whole-genome sequence variant databases

We used whole-genome sequencing (WGS) data of 139 pigs that were sequenced at an average read depth greater than 8x. Sequence reads were processed and aligned against the Sscrofa11.1 reference sequence as detailed in (Nosková et al., 2021). We called variants with DeepVariant (version 1.3.0, (Poplin et al., 2018; Yun et al., 2021)), producing a gVCF file per sample. The gVCF files were then merged and filtered using GLnexus (version 1.4.1, (Lin et al., n.d.)) with the DeepVariantWGS configuration, followed by imputation with Beagle 4.1 (Browning & Browning, 2007). Sequence variants were called previously for 266 cattle, 161 sheep and 157 goats (Table 1). We considered only biallelic sequence variants for our analyses.

### Public variant databases

We downloaded reference sequences and their annotations including non-coding RNAs, as well as a VCF file with polymorphic positions for 30 species from EVA (release 4, (Cezard et al., 2022)), Ensembl (release 107, (Cunningham et al., 2022)), or Ensembl Plants (release 55, (Bolser et al., 2016)). Access information for all data is provided in Table S2.

We used these data to evaluate the number of variants, proportion of variants in protein-coding regions, average genome-wide variability (in variants per 100 bases) and transition to transversion (Ti/Tv) ratio. Variants overlapping exons, start codons and stop codons were considered as coding variants.

### Prediction of branch point sequences

We followed the approach of Kadri et al. (Kadri et al., 2021) to predict BPS in 30 species using BPP (Q. Zhang et al., 2017). In short, we obtained coordinates of introns in protein-coding genes from GTF files of each species, mindful of gene-strand orientation. We used species-specific weighted octanucleotide frequencies estimated as suggested by Zhang et al. (Q. Zhang et al., 2017) and the position weight matrix of predicted human BPS for model training (Q. Zhang et al., 2017). For the analysis of constraint, we only considered the most probable BPS within each intron.

The variability between positions 4 and 6 of the heptamer was compared for each species with Fisher’s exact test. We applied Bonferroni-correction to account for multiple testing (number of species tested).

### Variation in genic features

Variability was calculated as the number of variants per 100 bp divided by the respective species’ genome-wide variability for variants overlapping nine annotated genomic features (3’ and 5’ splice sites, start and stop codons, 3’ and 5’ UTR, introns, exons, intergenic regions) and predicted BPS. Genome-wide variability, i.e., average number of variants per 100 bp, was calculated as total number of variants divided by the size of the genome. The genome size was the total length of all chromosomes considered but undetermined bases (“N”) were excluded.

### Genomic variant database analysis

Based on the analyses of WGS datasets we established three criteria to assess the quality of public databases. The criteria were (i) genome-wide variability of minimum 1 variable site per 1000 bp; (ii) variability in intergenic regions above the average genome-wide; (iii) depletion of variation at the 4 bases overlapping splice sites.

Databases not fulfilling all criteria were excluded from further analyses (Table S1, File S1). Eleven species that satisfied these criteria were considered to estimate constraint patterns.

## Data Availability

Sequencing reads of 139 pigs are available at the European Nucleotide Archive (ENA) (http://www.ebi.ac.uk/ena) at BioProjects PRJEB38156, PRJEB37956, and PRJEB39374.

## References

Blakes, A. J. M., Wai, H. A., Davies, I., Moledina, H. E., Ruiz, A., Thomas, T., Bunyan, D., Thomas, N. S., Burren, C. P., Greenhalgh, L., Lees, M., Pichini, A., Smithson, S. F., Taylor Tavares, A. L., O’Donovan, P., Douglas, A. G. L., Whiffin, N., Baralle, D., & Lord, J. (2022). A systematic analysis of splicing variants identifies new diagnoses in the 100,000 Genomes Project. Genome Medicine, 14(1). https://doi.org/10.1186/S13073-022-01087-X

Bolser, D., Staines, D. M., Pritchard, E., & Kersey, P. (2016). Ensembl Plants: Integrating Tools for Visualizing, Mining, and Analyzing Plant Genomics Data. Methods in Molecular Biology (Clifton, N.J.), 1374, 115–140. https://doi.org/10.1007/978-1-4939-3167-5_6

Browning, S. R., & Browning, B. L. (2007). Rapid and accurate haplotype phasing and missing-data inference for whole-genome association studies by use of localized haplotype clustering. American Journal of Human Genetics, 81(5), 1084–1097. https://doi.org/10.1086/521987

Cezard, T., Cunningham, F., Hunt, S. E., Koylass, B., Kumar, N., Saunders, G., Shen, A., Silva, A. F., Tsukanov, K., Venkataraman, S., Flicek, P., Parkinson, H., & Keane, T. M. (2022). The European Variation Archive: A FAIR resource of genomic variation for all species. Nucleic Acids Research, 50(D1), D1216–D1220. https://doi.org/10.1093/nar/gkab960

Cunningham, F., Allen, J. E., Allen, J., Alvarez-Jarreta, J., Amode, M. R., Armean, I. M., Austine-Orimoloye, O., Azov, A. G., Barnes, I., Bennett, R., Berry, A., Bhai, J., Bignell, A., Billis, K., Boddu, S., Brooks, L., Charkhchi, M., Cummins, C., Da Rin Fioretto, L., … Flicek, P. (2022). Ensembl 2022. Nucleic Acids Research, 50(D1), D988–D995. https://doi.org/10.1093/NAR/GKAB1049

Gao, K., Masuda, A., Matsuura, T., & Ohno, K. (2008). Human branch point consensus sequence is yUnAy. Nucleic Acids Research, 36(7), 2257. https://doi.org/10.1093/NAR/GKN073

Johnston, J. J., & Biesecker, L. G. (2013). Databases of genomic variation and phenotypes: existing resources and future needs. Human Molecular Genetics, 22(R1), R27–R31. https://doi.org/10.1093/HMG/DDT384

Kadri, N. K., Mapel, X. M., & Pausch, H. (2021). The intronic branch point sequence is under strong evolutionary constraint in the bovine and human genome. Communications Biology 2021 4:1, 4(1), 1–13. https://doi.org/10.1038/s42003-021-02725-7

Keller, E. B., & Noon, W. A. (1984). Intron splicing: a conserved internal signal in introns of animal pre-mRNAs. Proceedings of the National Academy of Sciences of the United States of America, 81(23), 7417. https://doi.org/10.1073/PNAS.81.23.7417

Khafizov, K., Ivanov, M. V., Glazova, O. V., & Kovalenko, S. P. (2015). Computational approaches to study the effects of small genomic variations. Journal of Molecular Modeling 2015 21:10, 21(10), 1–14. https://doi.org/10.1007/S00894-015-2794-Y

Královičová, J., Lei, H., & Vořechovský, I. (2006). Phenotypic consequences of branch point substitutions. Human Mutation, 27(8), 803–813. https://doi.org/10.1002/HUMU.20362

LaDuca, H., Farwell, K. D., Vuong, H., Lu, H. M., Mu, W., Shahmirzadi, L., Tang, S., Chen, J., Bhide, S., & Chao, E. C. (2017). Exome sequencing covers >98% of mutations identified on targeted next generation sequencing panels. PloS One, 12(2). https://doi.org/10.1371/JOURNAL.PONE.0170843

Lee, Y., & Rio, D. C. (2015). Mechanisms and Regulation of Alternative Pre-mRNA Splicing. Annual Review of Biochemistry, 84, 291. https://doi.org/10.1146/ANNUREV-BIOCHEM-060614-034316

Li, C., Chen, B., Langda, S., Zhou, S., Kalds, P., Zhang, K., Bhati, M., Leonard, A., Zhu, X., Huang, S., Li, R., Cuoji, A., Wu, Y., Cuomu, R., Gui, B., Li, M., Wang, Y., Li, Y., Fang, W., … Wang, X. (2023). Comparative genomic analyses shed light on the genetic control of high-altitude adaptation in sheep. Submitted.

Li, C., Wu, Y., Chen, B., Cai, Y., Guo, J., Leonard, A. S., Kalds, P., Zhou, S., Zhang, J., Zhou, P., Gan, S., Jia, T., Pu, T., Suo, L., Li, Y., Zhang, K., Li, L., Purevdorj, M., Wang, X., … Wang, X. (2022). Markhor-derived Introgression of a Genomic Region Encompassing PAPSS2 Confers High-altitude Adaptability in Tibetan Goats. Molecular Biology and Evolution, 39(12). https://doi.org/10.1093/MOLBEV/MSAC253

Lin, M. F., Dnanexus, O. R., Penn, J., Bai, X., Reid, J. G., Krasheninina, O., & Salerno, W. J. (n.d.). GLnexus: joint variant calling for large cohort sequencing. https://doi.org/10.1101/343970

Maffucci, P., Bigio, B., Rapaport, F., Cobat, A., Borghesi, A., Lopez, M., Patin, E., Bolze, A., Shang, L., Bendavid, M., Scott, E. M., Stenson, P. D., Cunningham-Rundles, C., Cooper, D. N., Gleeson, J. G., Fellay, J., Quintana-Murci, L., Casanova, J. L., Abel, L., … Itan, Y. (2019). Blacklisting variants common in private cohorts but not in public databases optimizes human exome analysis. Proceedings of the National Academy of Sciences of the United States of America, 116(3), 950–959. https://doi.org/10.1073/PNAS.1808403116

Mercer, T. R., Clark, M. B., Andersen, S. B., Brunck, M. E., Haerty, W., Crawford, J., Taft, R. J., Nielsen, L. K., Dinger, M. E., & Mattick, J. S. (2015). Genome-wide discovery of human splicing branchpoints. Genome Research, 25(2), 290. https://doi.org/10.1101/GR.182899.114

Mitchell, A. A., Zwick, M. E., Chakravarti, A., & Cutler, D. J. (2004). Discrepancies in dbSNP confirmation rates and allele frequency distributions from varying genotyping error rates and patterns. Bioinformatics, 20(7), 1022–1032. https://doi.org/10.1093/BIOINFORMATICS/BTH034

Musumeci, L., Arthur, J. W., Cheung, F. S. G., Hoque, A., Lippman, S., & Reichardt, J. K. V. (2010). Single Nucleotide Differences (SNDs) in the dbSNP Database May Lead to Errors in Genotyping and Haplotyping Studies. Human Mutation, 31(1), 67. https://doi.org/10.1002/HUMU.21137

Nosková, A., Bhati, M., Kadri, N. K., Crysnanto, D., Neuenschwander, S., Hofer, A., & Pausch, H. (2021). Characterization of a haplotype-reference panel for genotyping by low-pass sequencing in Swiss Large White pigs. BMC Genomics, 22(1), 290. https://doi.org/10.1186/s12864-021-07610-5

Paggi, J. M., & Bejerano, G. (2018). A sequence-based, deep learning model accurately predicts RNA splicing branchpoints. RNA (New York, N.Y.), 24(12), 1647–1653. https://doi.org/10.1261/RNA.066290.118

Pineda, J. M. B., & Bradley, R. K. (2018). Most human introns are recognized via multiple and tissue-specific branchpoints. Genes and Development, 32(7–8), 577–591. https://doi.org/10.1101/GAD.312058.118

Poplin, R., Chang, P. C., Alexander, D., Schwartz, S., Colthurst, T., Ku, A., Newburger, D., Dijamco, J., Nguyen, N., Afshar, P. T., Gross, S. S., Dorfman, L., McLean, C. Y., & Depristo, M. A. (2018). A universal SNP and small-indel variant caller using deep neural networks. Nature Biotechnology 2018 36:10, 36(10), 983–987. https://doi.org/10.1038/nbt.4235

Schwartz, S. H., Silva, J., Burstein, D., Pupko, T., Eyras, E., & Ast, G. (2008). Large-scale comparative analysis of splicing signals and their corresponding splicing factors in eukaryotes. Genome Research, 18(1), 88. https://doi.org/10.1101/GR.6818908

Signal, B., Gloss, B. S., Dinger, M. E., & Mercer, T. R. (2018). Machine learning annotation of human branchpoints. Bioinformatics (Oxford, England), 34(6), 920–927. https://doi.org/10.1093/BIOINFORMATICS/BTX688

Taggart, A. J., Lin, C. L., Shrestha, B., Heintzelman, C., Kim, S., & Fairbrother, W. G. (2017). Large-scale analysis of branchpoint usage across species and cell lines. Genome Research, 27(4), 639–649. https://doi.org/10.1101/GR.202820.115/-/DC1

Yun, T., Li, H., Chang, P. C., Lin, M. F., Carroll, A., & McLean, C. Y. (2021). Accurate, scalable cohort variant calls using DeepVariant and GLnexus. Bioinformatics, 36(24), 5582–5589. https://doi.org/10.1093/BIOINFORMATICS/BTAA1081

Zhang, P., Philippot, Q., Ren, W., Lei, W. Te, Li, J., Stenson, P. D., Palacín, P. S., Colobran, R., Boisson, B., Zhang, S. Y., Puel, A., Pan-Hammarström, Q., Zhang, Q., Cooper, D. N., Abel, L., & Casanova, J. L. (2022). Genome-wide detection of human variants that disrupt intronic branchpoints. Proceedings of the National Academy of Sciences of the United States of America, 119(44), e2211194119. https://doi.org/10.1073/PNAS.2211194119/SUPPL_FILE/PNAS.2211194119.SD06.PDF

Zhang, Q., Fan, X., Wang, Y., Sun, M.-A., Shao, J., & Guo, D. (2017). BPP: a sequence-based algorithm for branch point prediction. Bioinformatics, 33(20), 3166–3172. https://doi.org/10.1093/bioinformatics/btx401

Zhang, X., Zhang, Y., Wang, T., Li, Z., Cheng, J., Ge, H., Tang, Q., Chen, K., Liu, L., Lu, C., Guo, J., Zheng, B., & Zheng, Y. (2019). A Comprehensive Map of Intron Branchpoints and Lariat RNAs in Plants. The Plant Cell, 31(5), 956. https://doi.org/10.1105/TPC.18.00711

Zhuang, Y., Goldstein, A. M., & Weiner, A. M. (1989). UACUAAC is the preferred branch site for mammalian mRNA splicing. Proceedings of the National Academy of Sciences of the United States of America, 86(8), 2752–2756. https://doi.org/10.1073/PNAS.86.8.2752

