## Supplementary Figures for "Exploiting public databases of genomic variation to quantify evolutionary constraint on the branch point sequence in 30 plant and animal species"

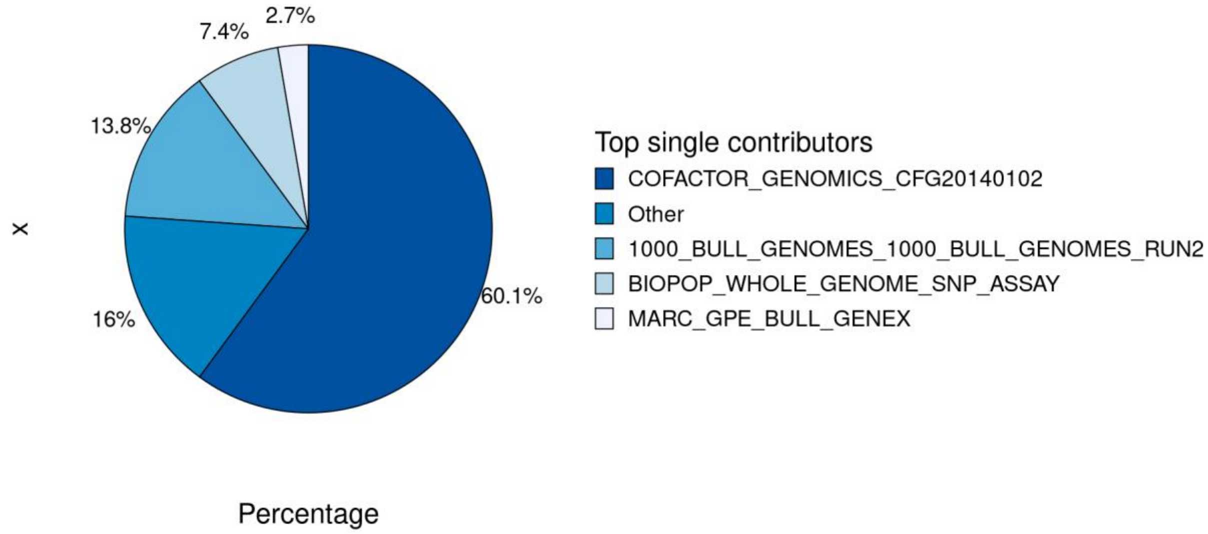


**Figure S1**: **Main contributors of single-entry variants.** Top four contributors submitted 84 % of all (N = 51,359,349) singletons. The largest batch (N = 31,580,941) of unvalidated variants was submitted by COFACTOR_GENOMICS_CFG20140112.

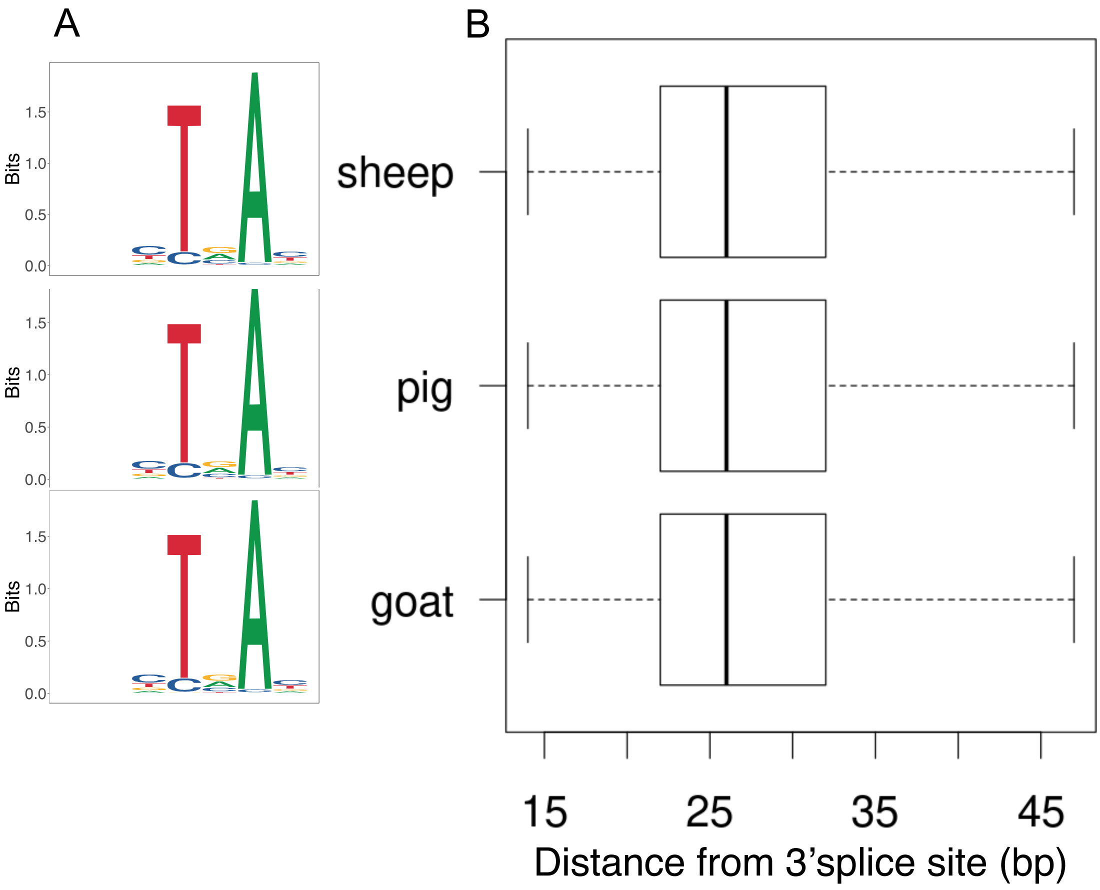


**Figure S2: Predicted branch point sequences in sheep, pig and goat.** (A) Motif logos of predicted branch point consensus sequences and (B) their placement (distance from 3' splice site in base pairs) in the sheep, pig and goat genomes.

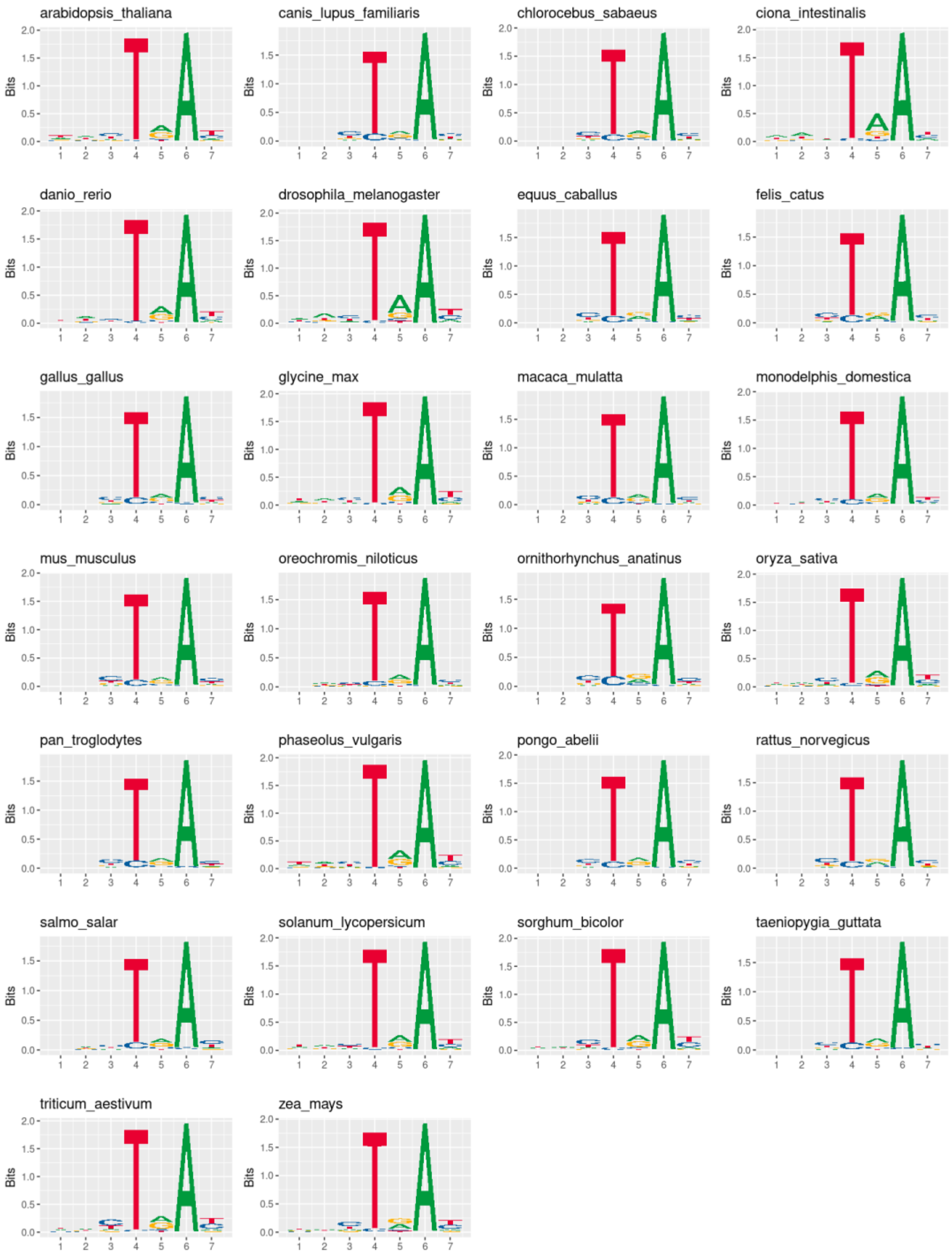


**Figure S3**: **Predicted branch point consensus sequences in 26 species.**

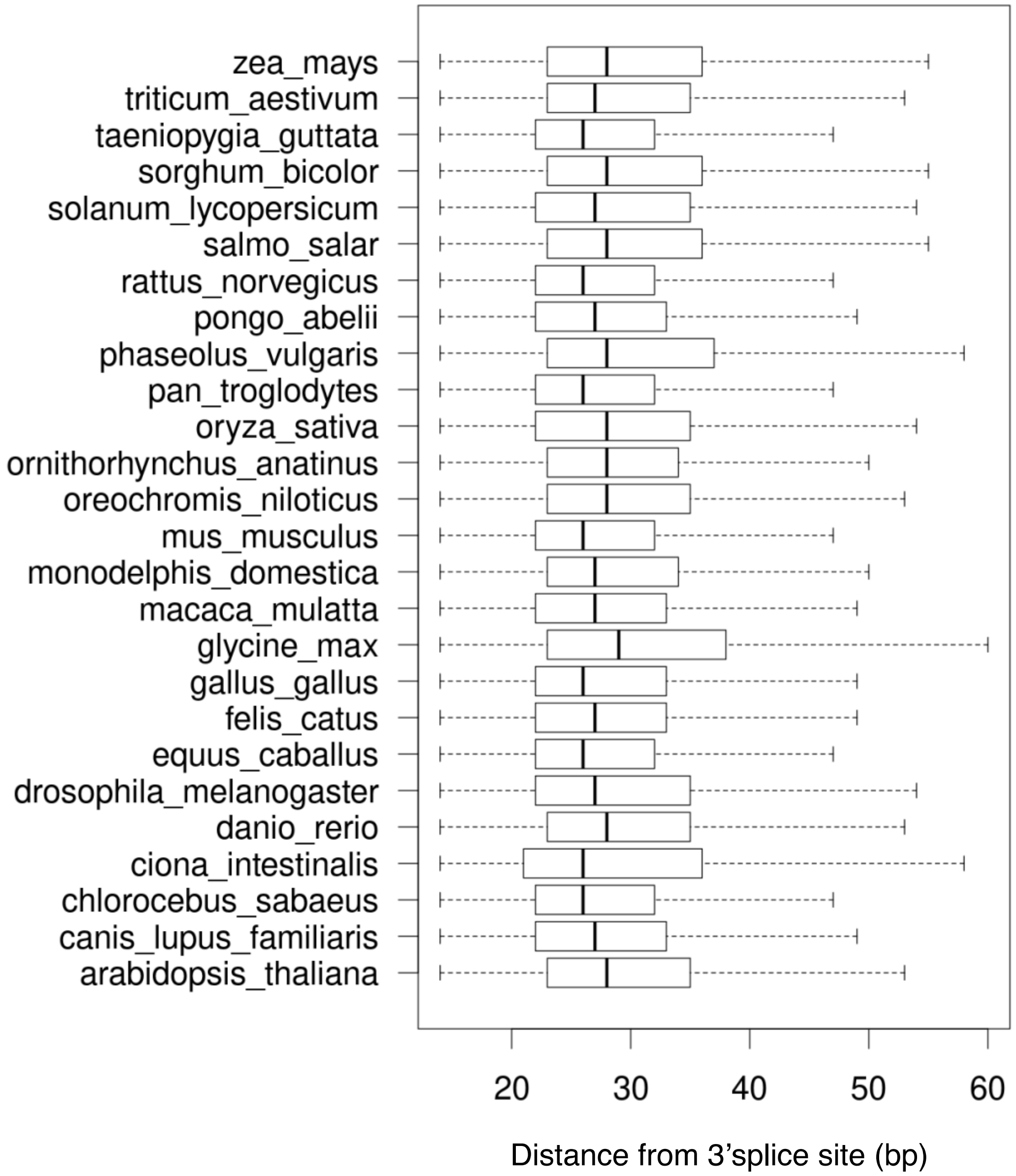


**Figure S4: Placement of predicted branch points in 26 species.**
