## Supplementary File1 for "Exploiting public databases of genomic variation to quantify evolutionary constraint on the branch point sequence in 30 plant and animal species"

**File S1**: Variability in nine genomic features, splice sites and predicted branch point sequences assessed from public variant databases in 26 species. Red line denotes genome-wide variability (unless the variability is above the plotted range). Blue lines in the middle and right-most plots denote exome variability. Databases for Eleven species (marked with asterisk) passed our criteria, while 15 species (marked with 'X') failed for following reasons: two species had too low genome-wide variability (*Monodelphis domestica, Ornithorhynchus anatinus)*, 12 species had intergenic variability lower than the genome-wide (*Canis lupus familiaris, Ciona intestinalis, Danio rerio, Drosophila melanogaster, Felis catus, Gallus gallus, Oreochromis niloticus, Ornithorhynchus anatinus, Salmo salar, Taeniopygia guttata, Triticum aestivum, Zea mays*), and five species revealed implausible constraint at the splice sites (*Equus caballus, Gallus gallus, Salmo salar, Sorghum bicolor and Triticum aestivum*).

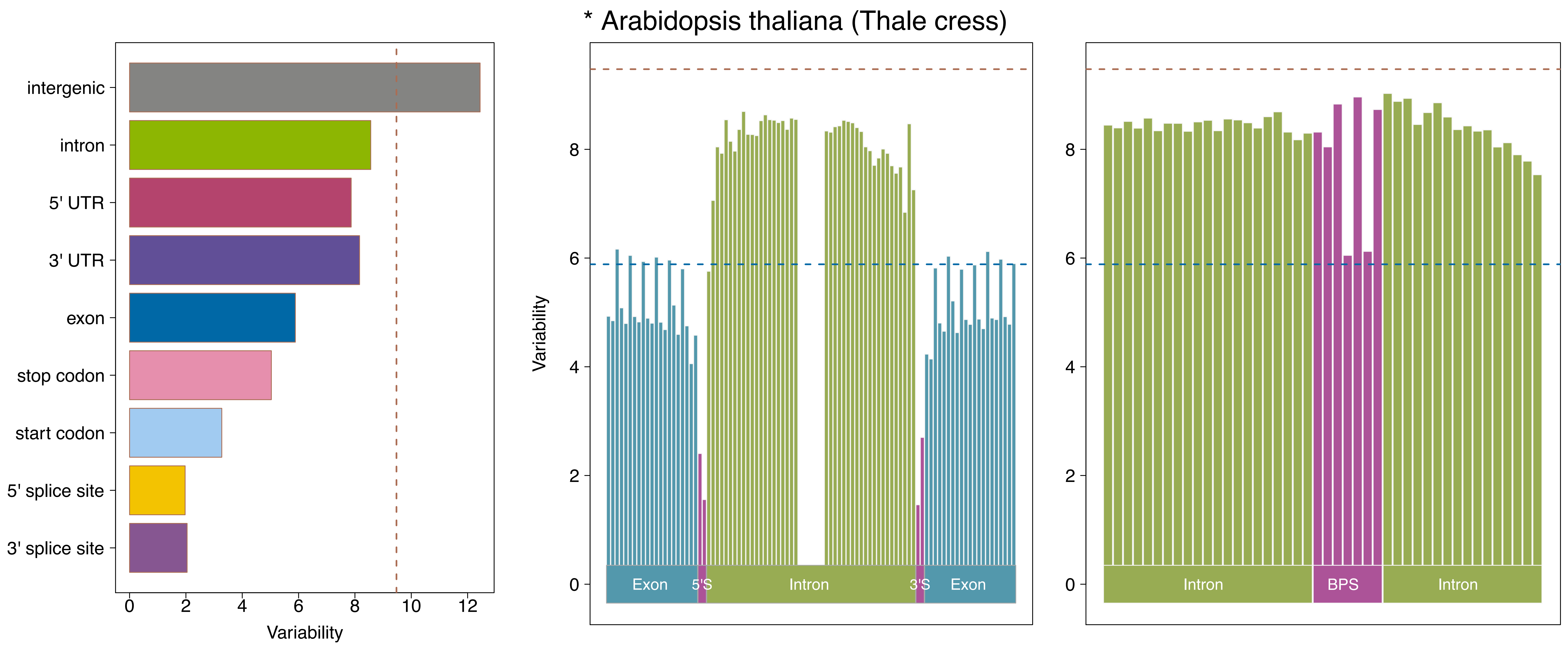

* Arabidopsis thaliana (Thale cress)
intergenic
intron
5' UTR
3' UTR
exon
stop codon
start codon
5' splice site
3' splice site
0
2
4
6
8
10
12
Variability
Variability
0
2
4
6
8
Exon
5'S
Intron
3'S
Exon
0
2
4
6
8
Intron
BPS
Intron

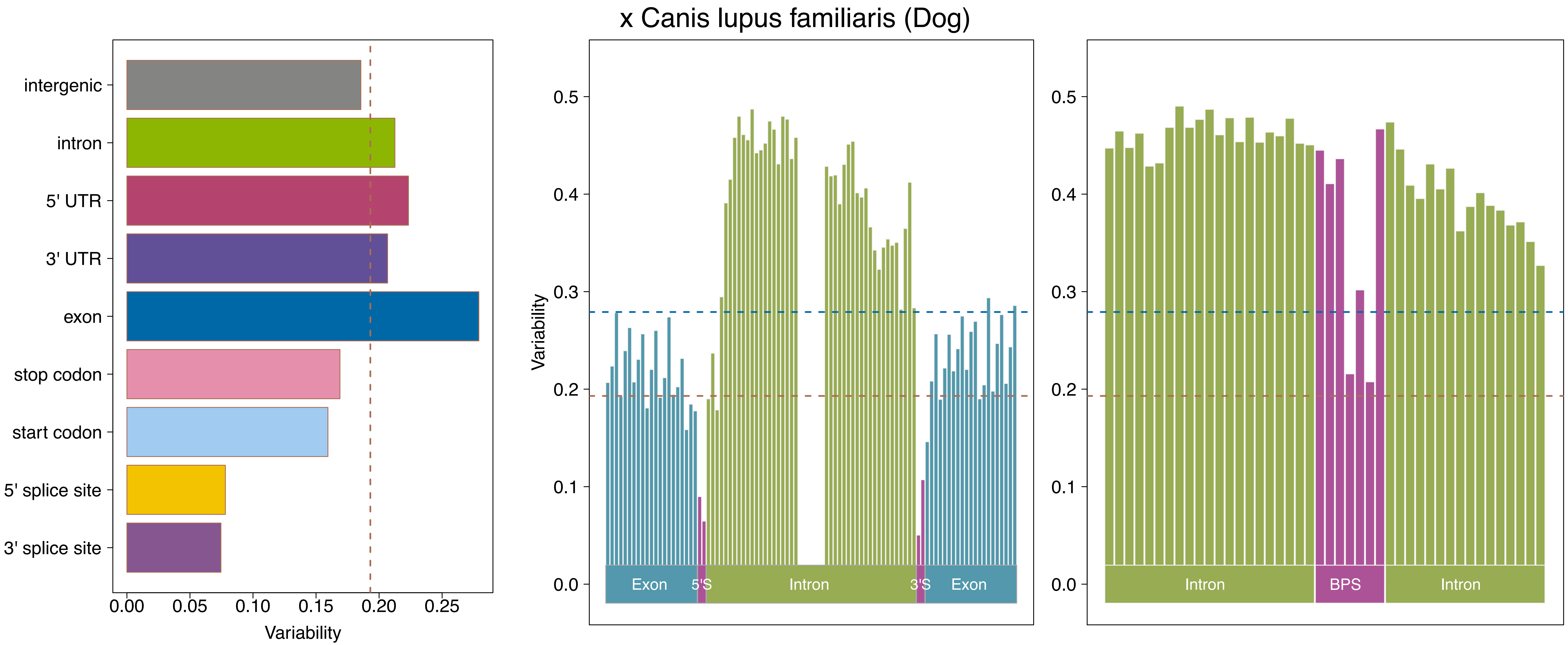

x Canis lupus familiaris (Dog)
intergenic
intron
5' UTR
3' UTR
exon
stop codon
start codon
5' splice site
3' splice site
0.00
0.05
0.10
0.15
0.20
0.25
Variability
Variability
0.0
0.1
0.2
0.3
0.4
0.5
Exon
5'S
Intron
3'S
Exon
0.0
0.1
0.2
0.3
0.4
0.5
Intron
BPS
Intron

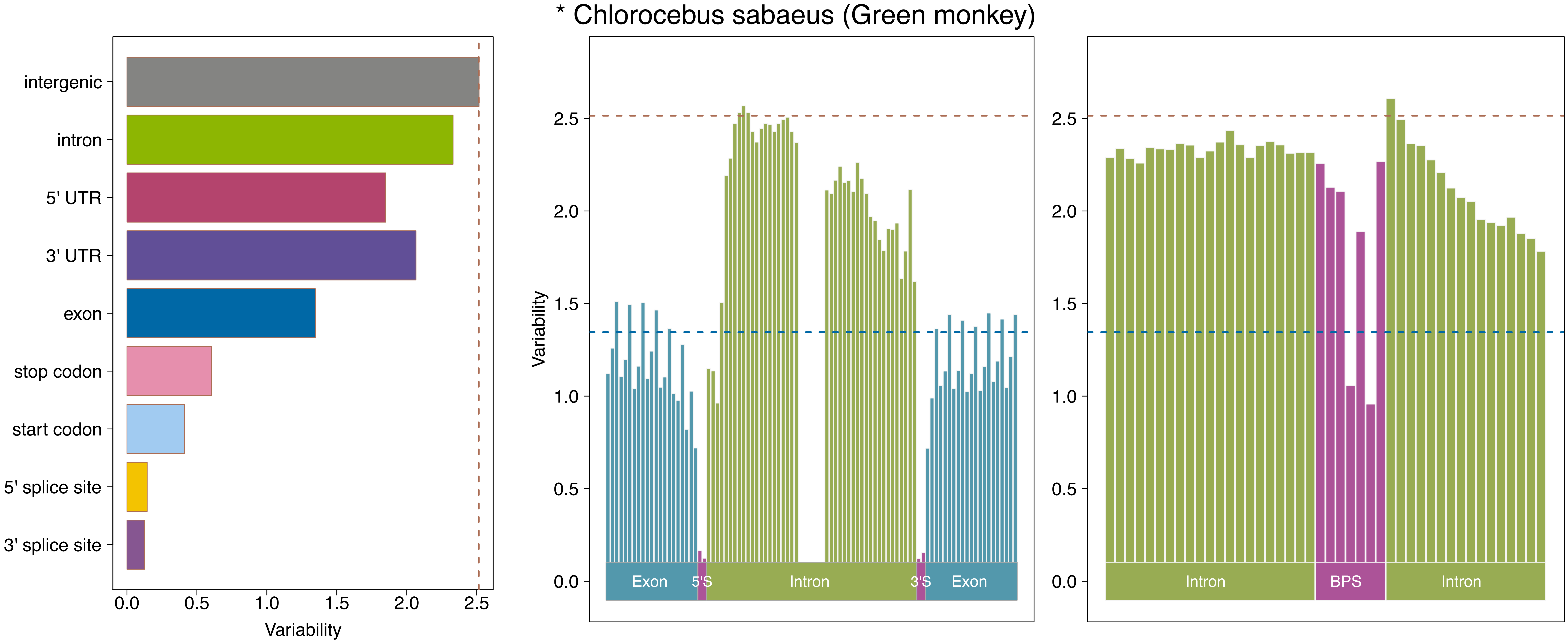
* Chlorocebus sabaeus (Green monkey)
intergenic
intron
5' UTR
3' UTR
exon
stop codon
start codon
5' splice site
3' splice site
0.0
0.5
1.0
1.5
2.0
2.5
Variability
Exon
5'S
Intron
3'S
Exon
Intron
BPS
Intron

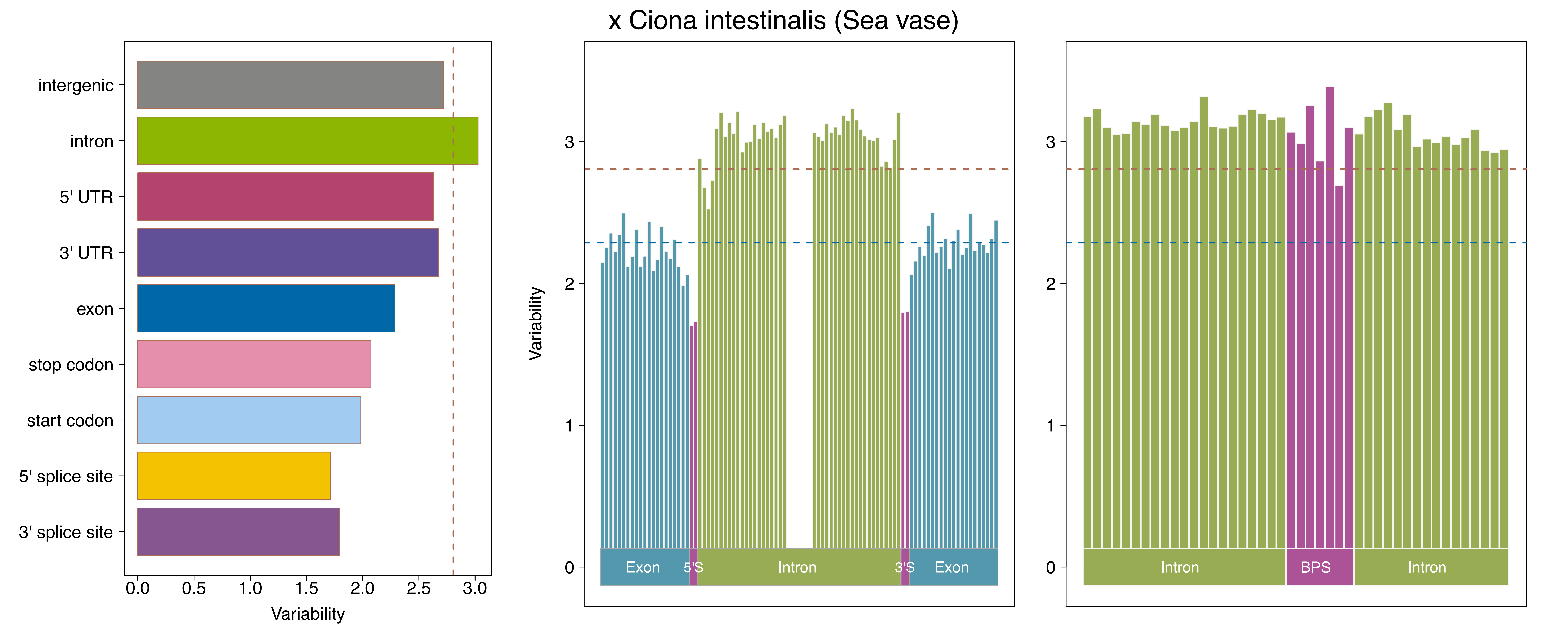
x Ciona intestinalis (Sea vase)
intergenic
intron
5' UTR
3' UTR
exon
stop codon
start codon
5' splice site
3' splice site
0.0
0.5
1.0
1.5
2.0
2.5
3.0
Variability
Variability
0
1
2
3
Exon
5'S
Intron
3'S
Exon
Intron
BPS
Intron

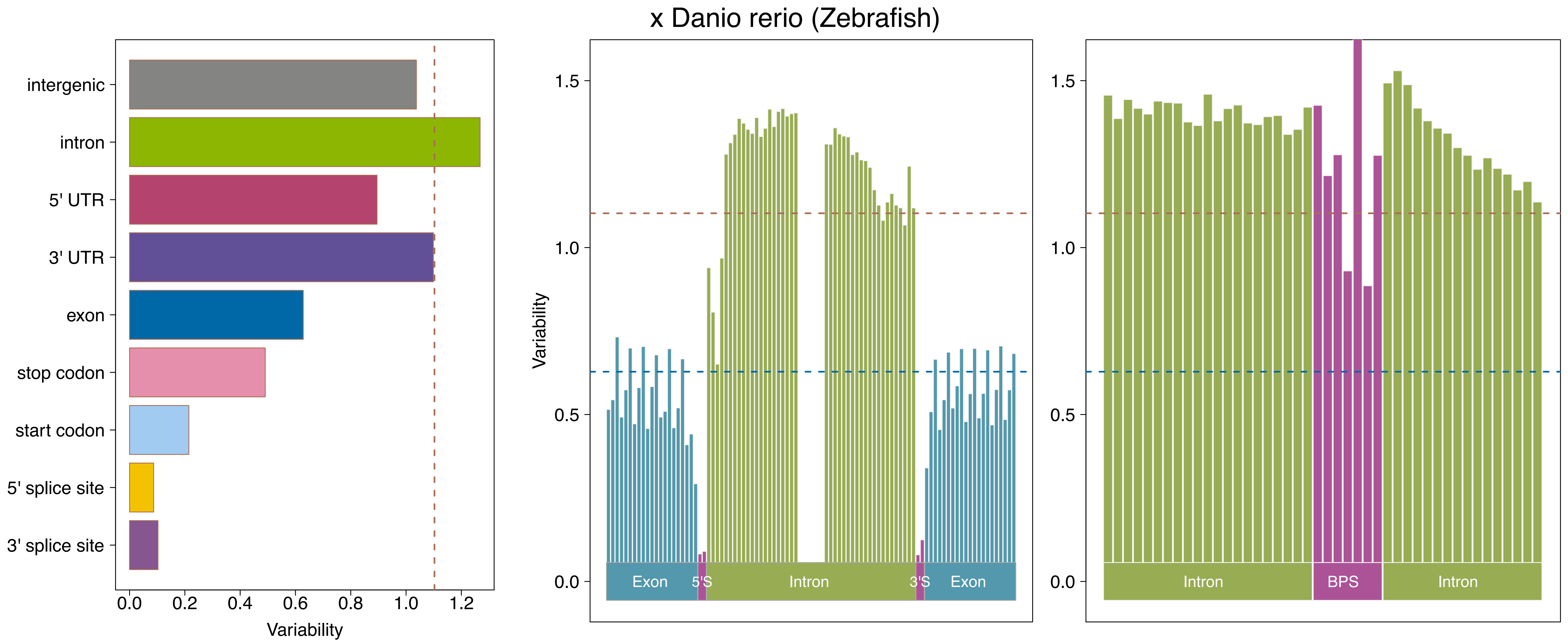

x Danio rerio (Zebrafish)
intergenic
intron
5' UTR
3' UTR
exon
stop codon
start codon
5' splice site
3' splice site
0.0
0.2
0.4
0.6
0.8
1.0
1.2
Variability
Variability
1.5
1.0
0.5
0.0
Exon
5'S
Intron
3'S
Exon
1.5
1.0
0.5
0.0
Intron
BPS
Intron

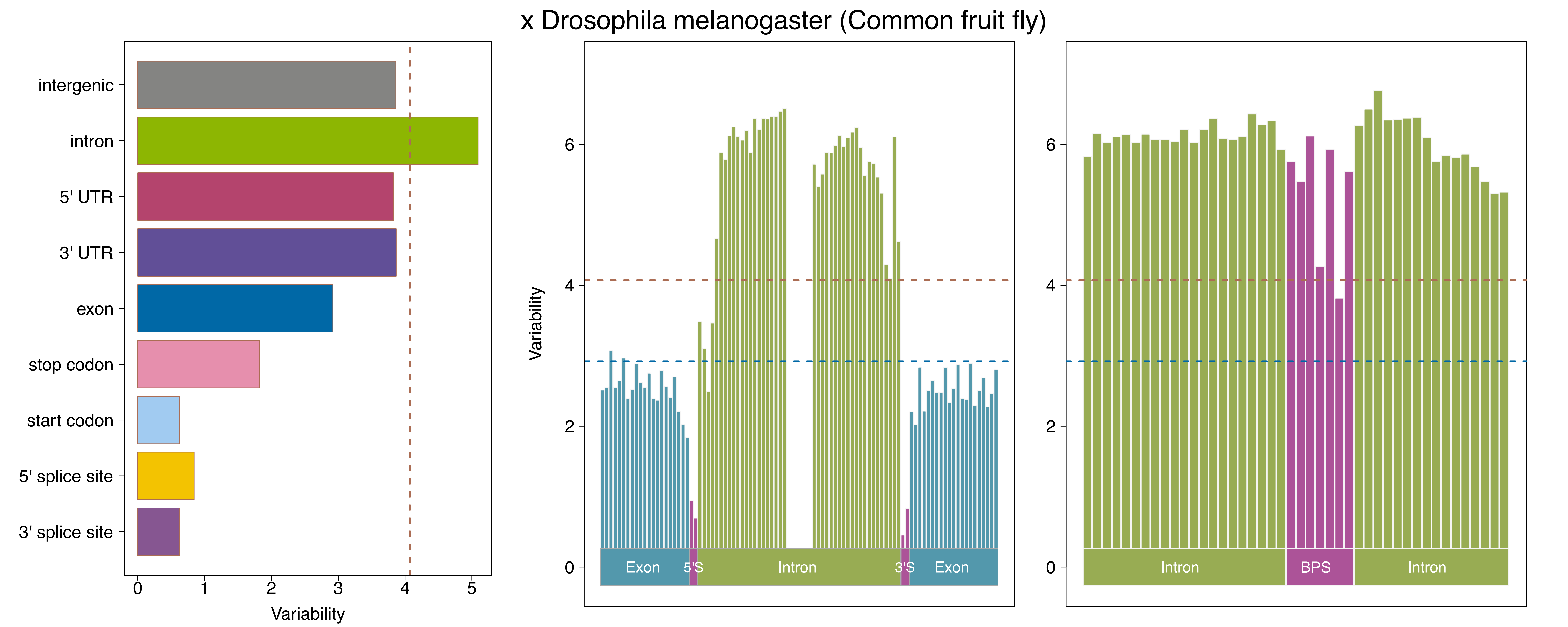
x Drosophila melanogaster (Common fruit fly)
intergenic
intron
5' UTR
3' UTR
exon
stop codon
start codon
5' splice site
3' splice site
0
1
2
3
4
5
Variability
Variability
0
2
4
6
Exon
5'S
Intron
3'S
Exon
0
2
4
6
Intron
BPS
Intron

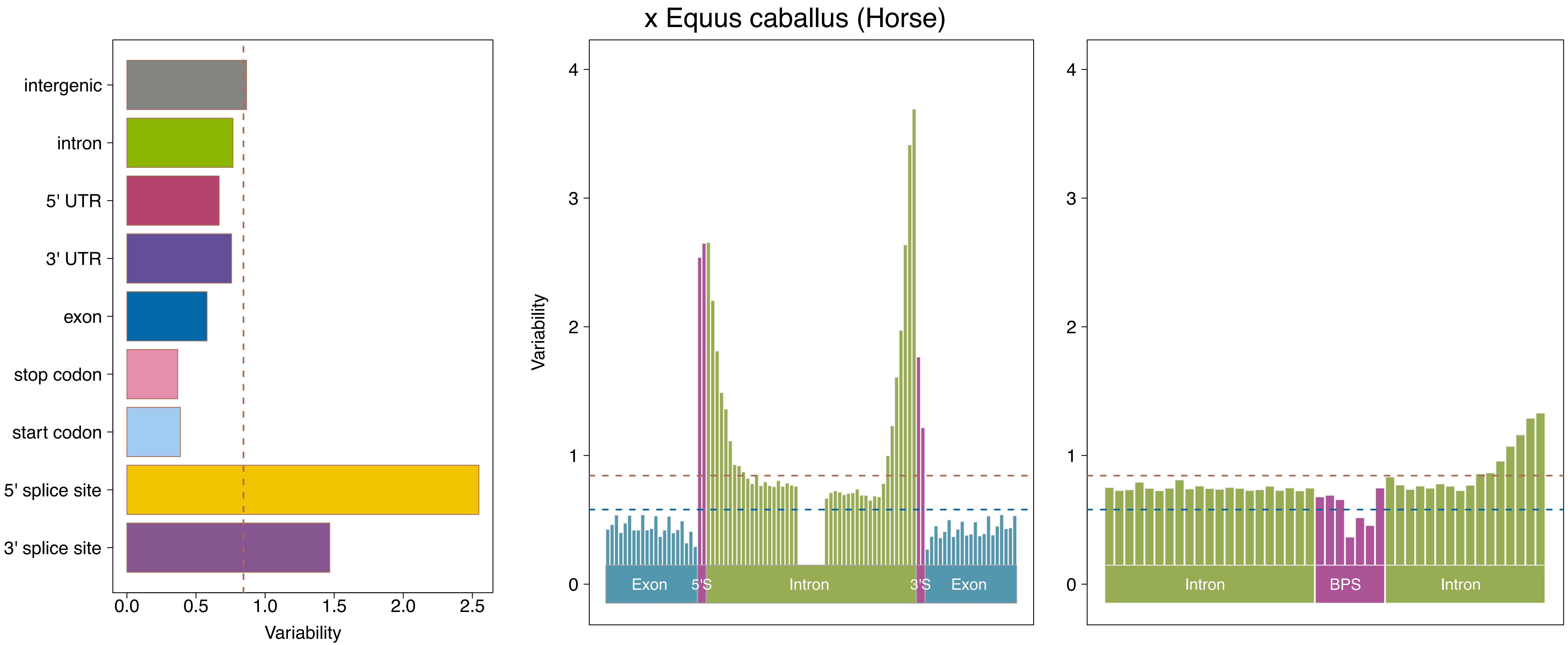

x Equus caballus (Horse)
intergenic
intron
5' UTR
3' UTR
exon
stop codon
start codon
5' splice site
3' splice site
0.0
0.5
1.0
1.5
2.0
2.5
Variability
Variability
0
1
2
3
4
Exon
5'S
Intron
3'S
Exon
0
1
2
3
4
Intron
BPS
Intron

x Felis catus (Domestic cat)

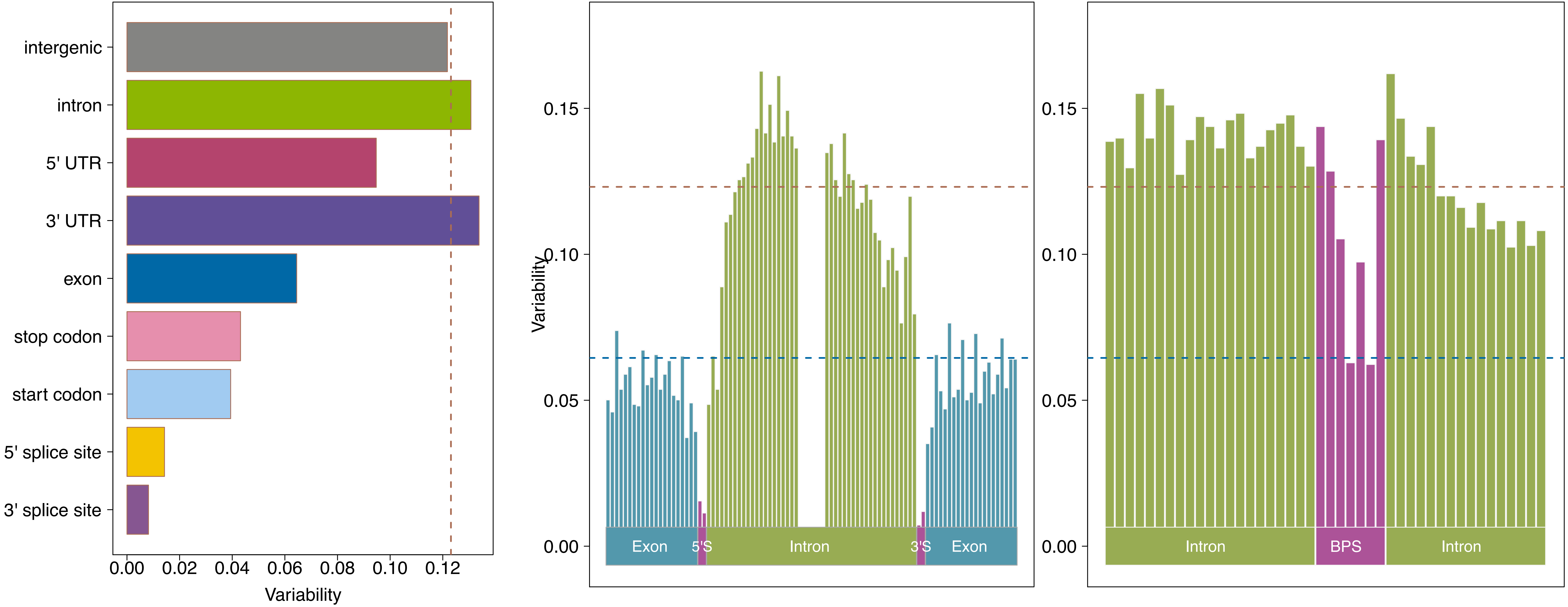

### x Gallus gallus (Chicken)

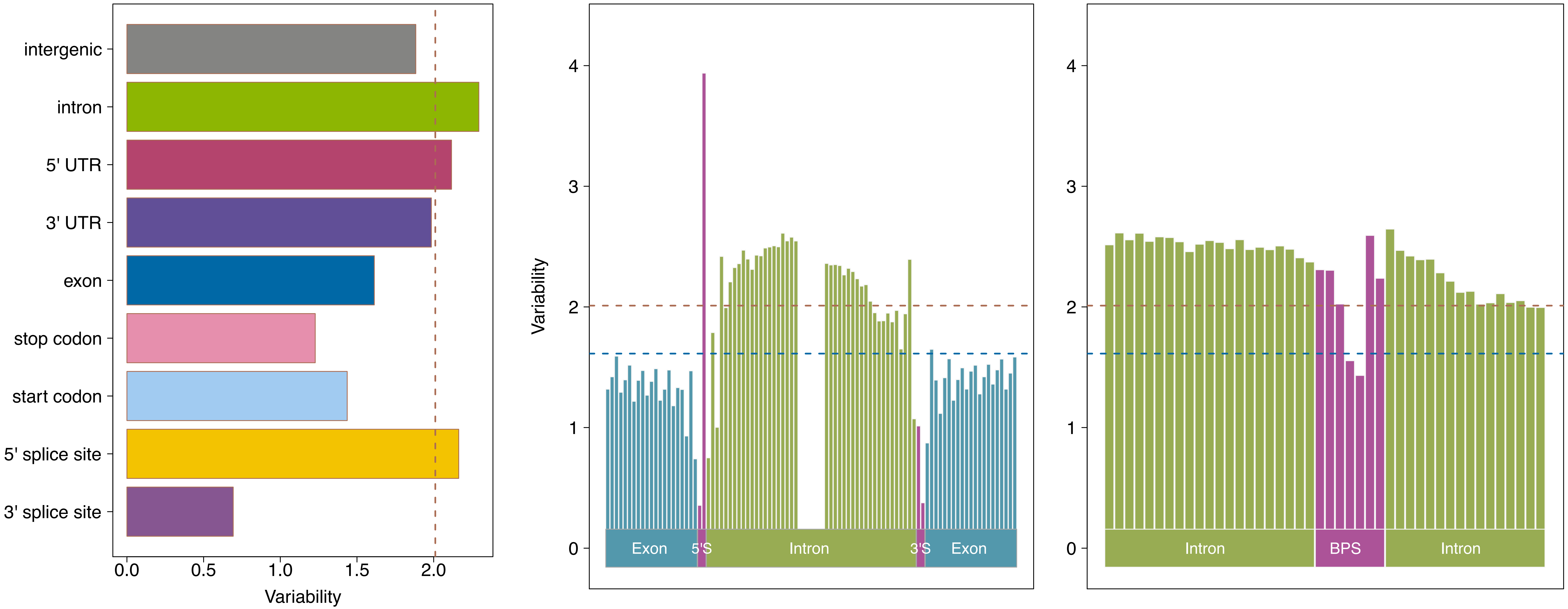

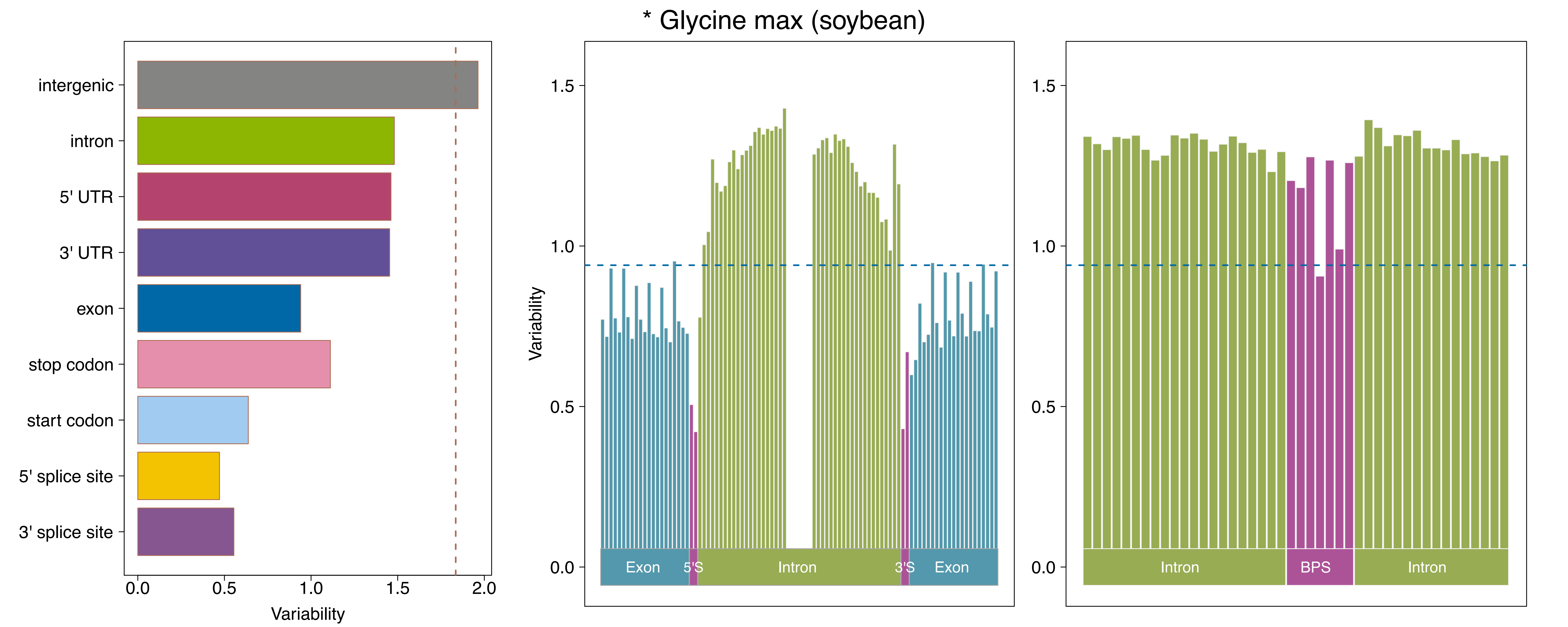
* Glycine max (soybean)
intergenic
intron
5' UTR
3' UTR
exon
stop codon
start codon
5' splice site
3' splice site
0.0
0.5
1.0
1.5
2.0
Variability
Exon
5'S
Intron
3'S
Exon
BPS
Intron
Intron
0.0
0.5
1.0
1.5
Variability

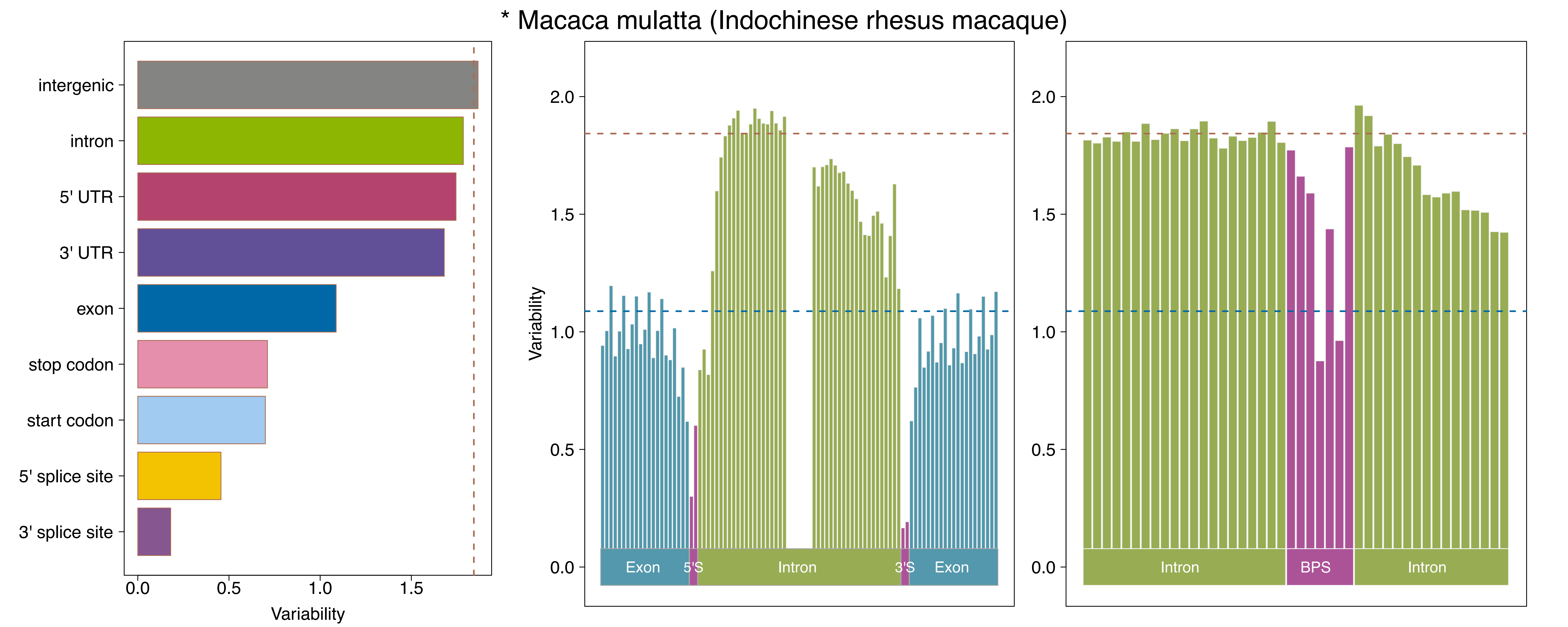
* Macaca mulatta (Indochinese rhesus macaque)
intergenic
intron
5' UTR
3' UTR
exon
stop codon
start codon
5' splice site
3' splice site
0.0
0.5
1.0
1.5
Variability
2.0
1.5
1.0
0.5
0.0
Variability
Exon
5'S
Intron
3'S
Exon
2.0
1.5
1.0
0.5
0.0
Intron
BPS
Intron

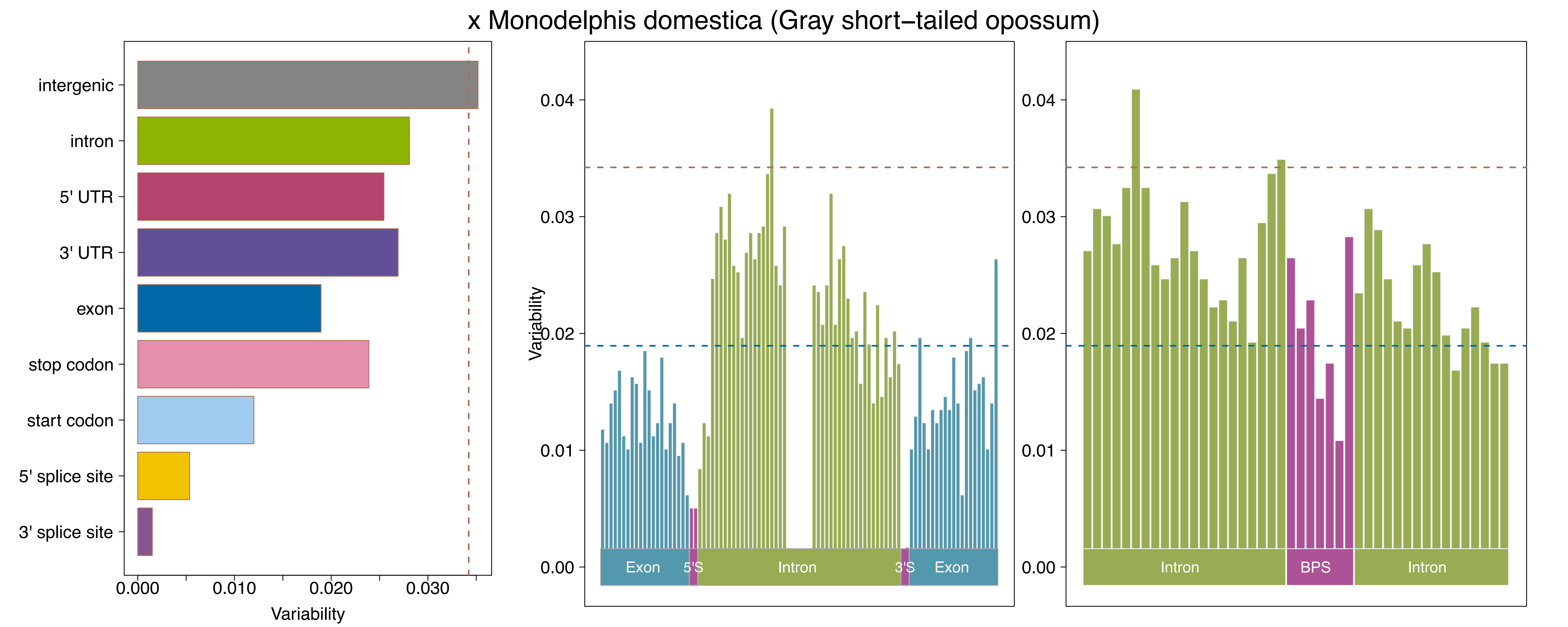

x Monodelphis domestica (Gray short–tailed opossum)
intergenic
intron
5' UTR
3' UTR
exon
stop codon
start codon
5' splice site
3' splice site
0.000
0.010
0.020
0.030
Variability
0.00
0.01
0.02
0.03
0.04
Variability
Exon
5'S
Intron
3'S
Exon
0.00
0.01
0.02
0.03
0.04
Intron
BPS
Intron

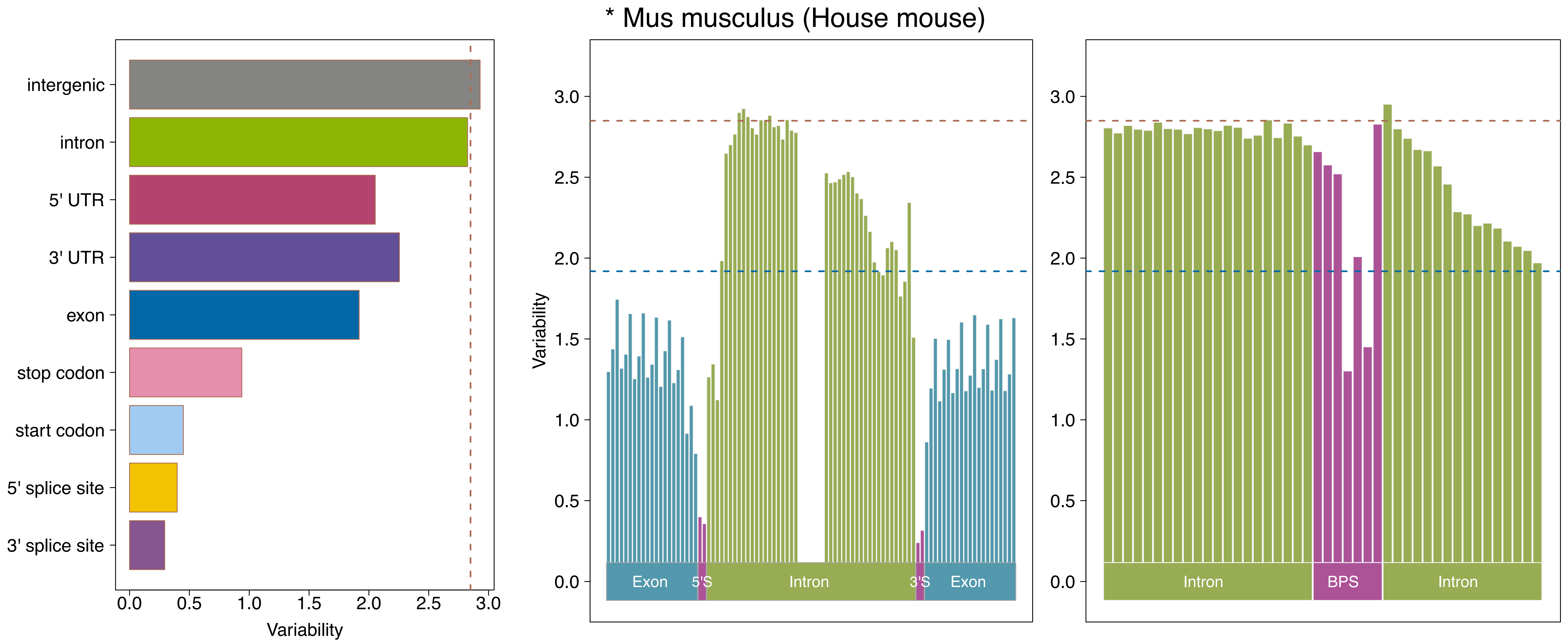

* Mus musculus (House mouse)
intergenic
intron
5' UTR
3' UTR
exon
stop codon
start codon
5' splice site
3' splice site
0.0
0.5
1.0
1.5
2.0
2.5
3.0
Variability
Variability
Exon
5'S
Intron
3'S
Exon
Intron
BPS
Intron

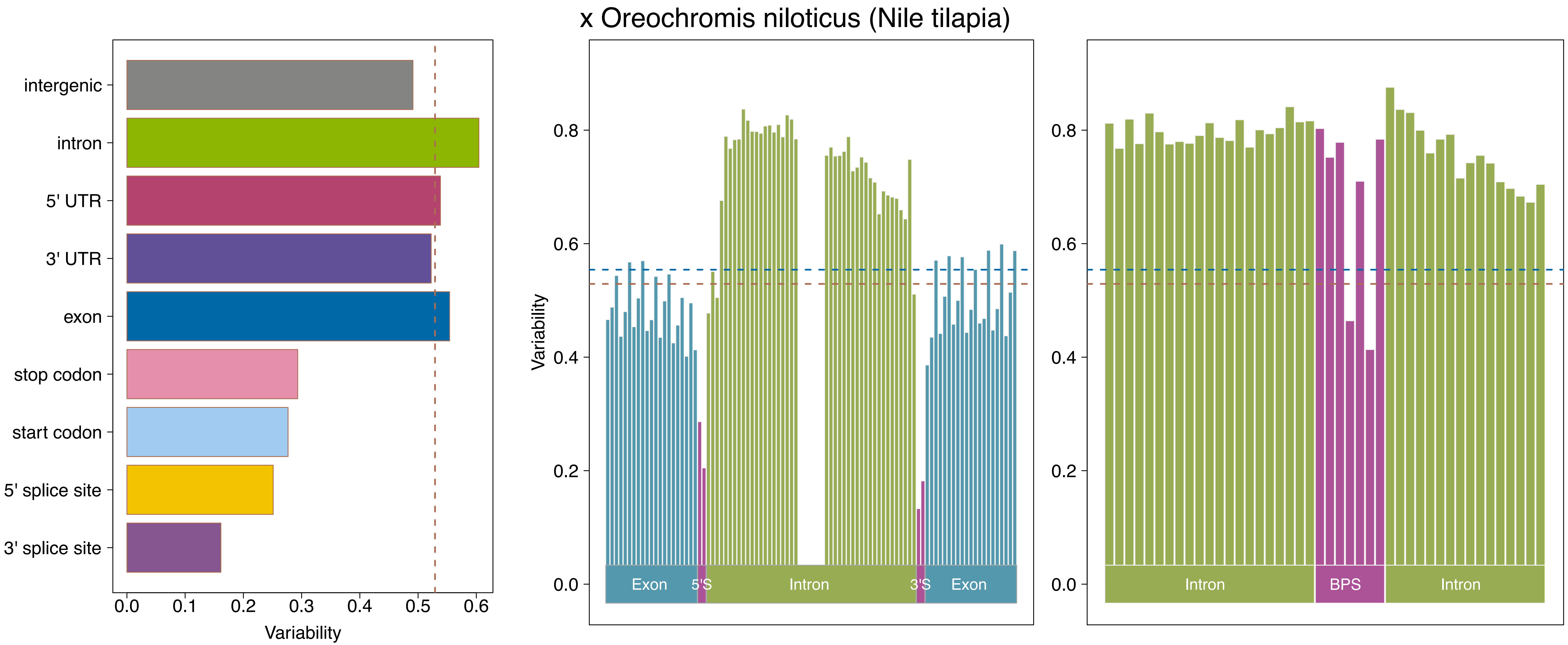

x Oreochromis niloticus (Nile tilapia)
intergenic
intron
5' UTR
3' UTR
exon
stop codon
start codon
5' splice site
3' splice site
0.0
0.1
0.2
0.3
0.4
0.5
0.6
Variability
Variability
0.0
0.2
0.4
0.6
0.8
Exon
5'S
Intron
3'S
Exon
0.0
0.2
0.4
0.6
0.8
Intron
BPS
Intron

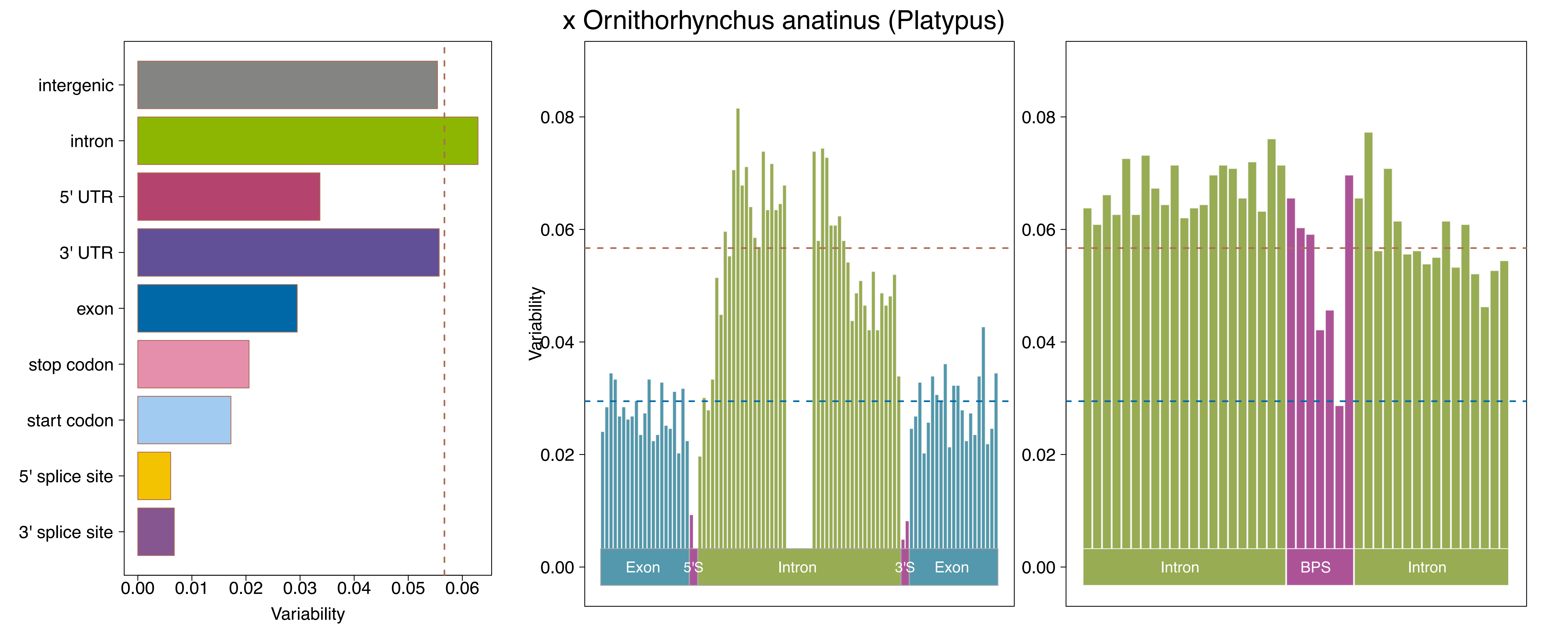

x Ornithorhynchus anatinus (Platypus)
intergenic
intron
5' UTR
3' UTR
exon
stop codon
start codon
5' splice site
3' splice site
0.00
0.01
0.02
0.03
0.04
0.05
0.06
Variability
0.08
0.06
0.04
0.02
0.00
Exon
5'S
Intron
3'S
Exon
Intron
BPS
Intron

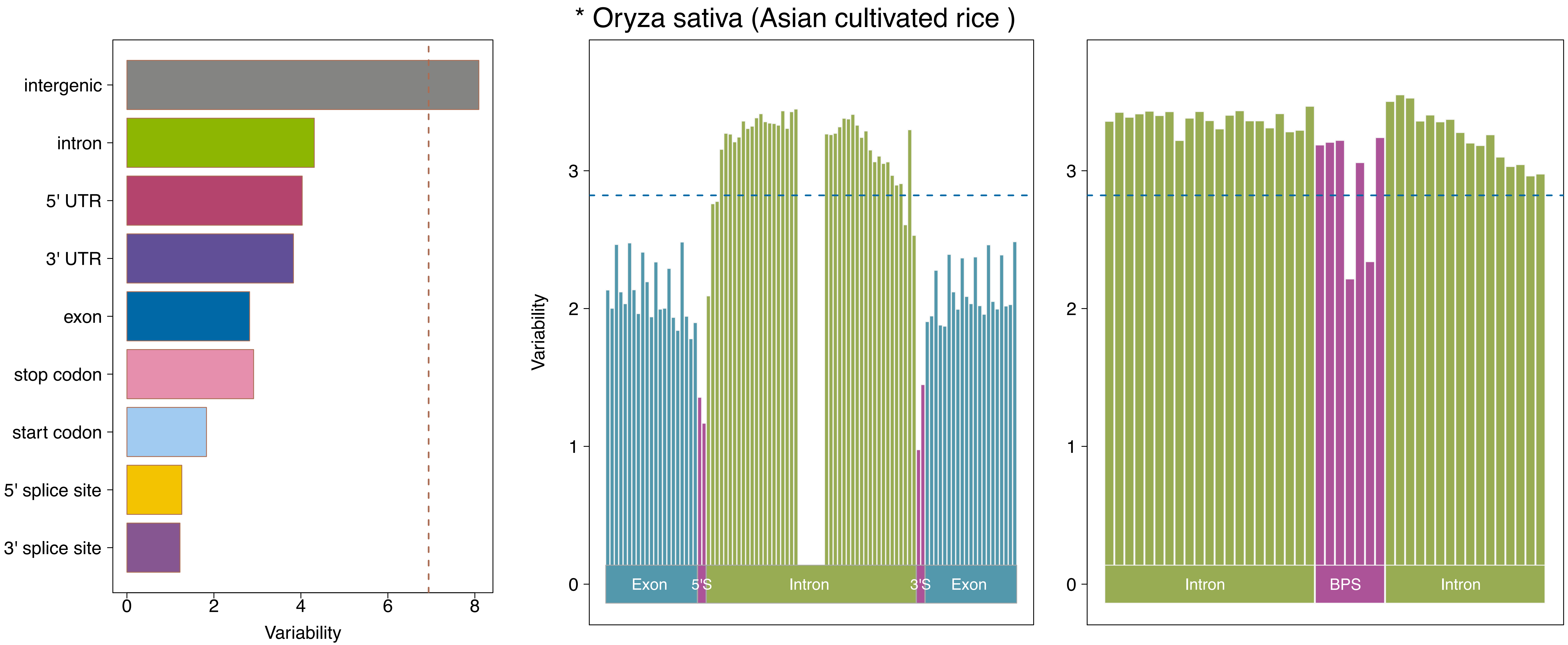
* Oryza sativa (Asian cultivated rice )
intergenic
intron
5' UTR
3' UTR
exon
stop codon
start codon
5' splice site
3' splice site
0
2
4
6
8
Variability
Variability
0
1
2
3
Exon
5'S
Intron
3'S
Exon
0
1
2
3
Intron
BPS
Intron

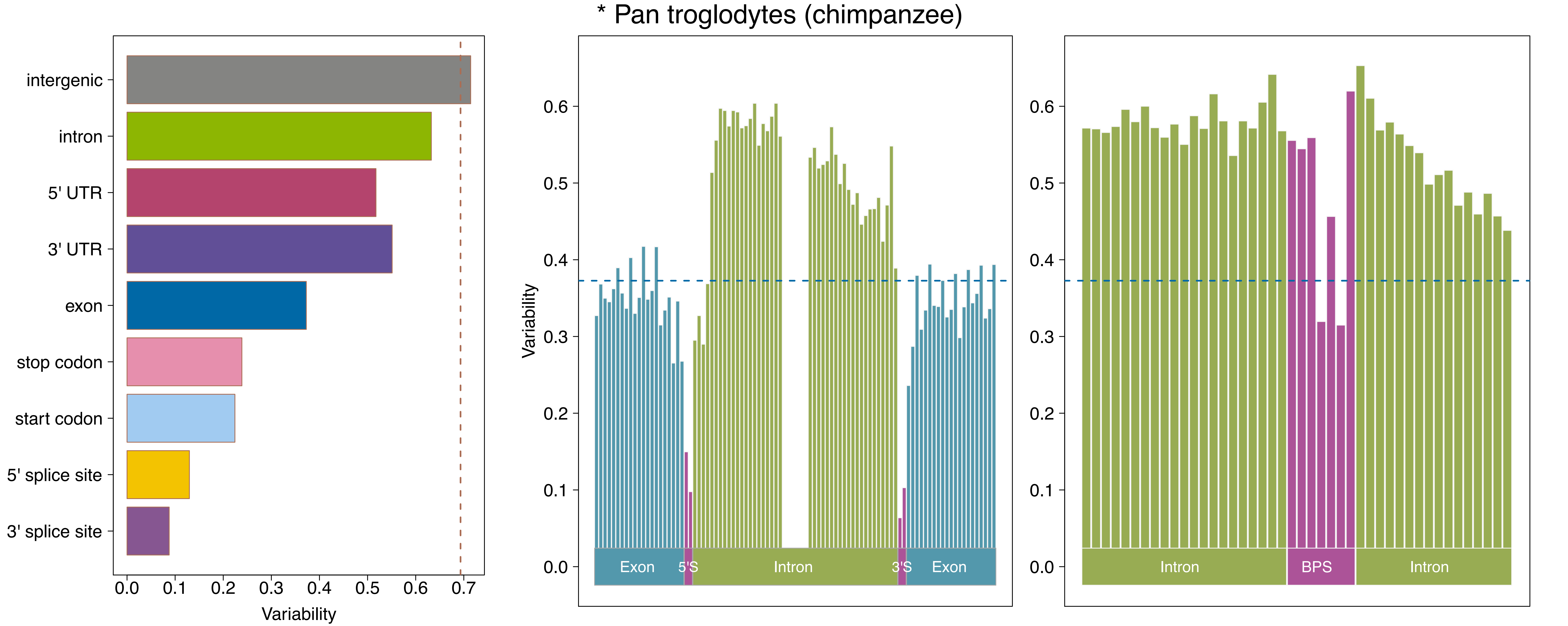
* Pan troglodytes (chimpanzee)
intergenic
intron
5' UTR
3' UTR
exon
stop codon
start codon
5' splice site
3' splice site
0.0
0.1
0.2
0.3
0.4
0.5
0.6
0.7
Variability
Variability
Exon
5'S
Intron
3'S
Exon
Intron
BPS
Intron

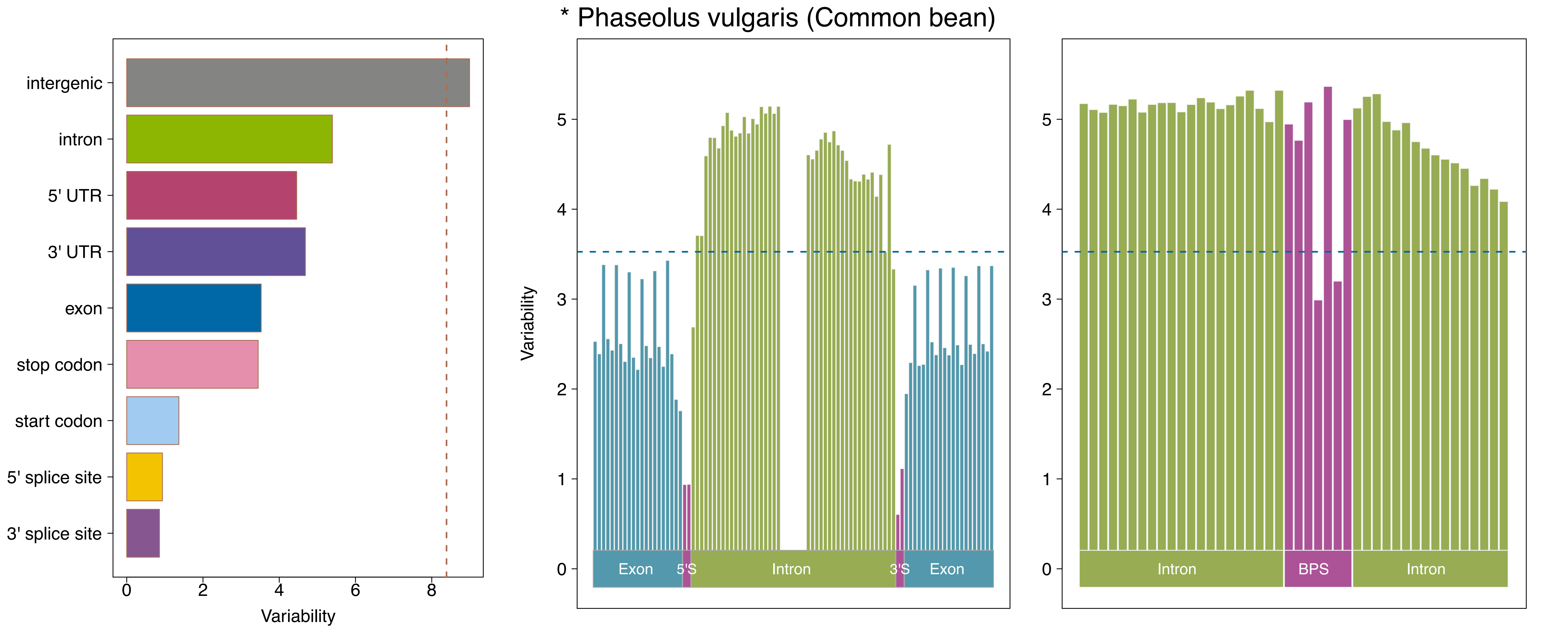
* Phaseolus vulgaris (Common bean)
intergenic
intron
5' UTR
3' UTR
exon
stop codon
start codon
5' splice site
3' splice site
0
2
4
6
8
Variability
Variability
0
1
2
3
4
5
Exon
5'S
Intron
3'S
Exon
0
1
2
3
4
5
Intron
BPS
Intron

* Pongo abelii (Sumatran orangutan)

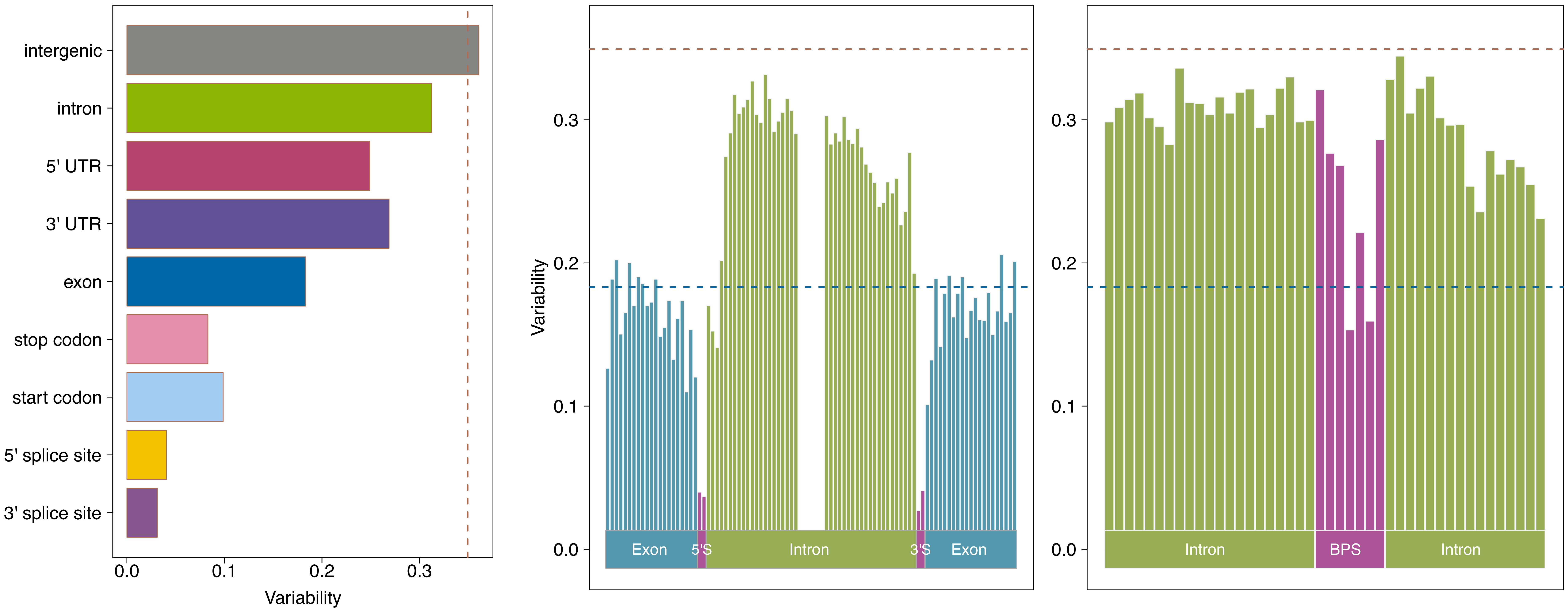

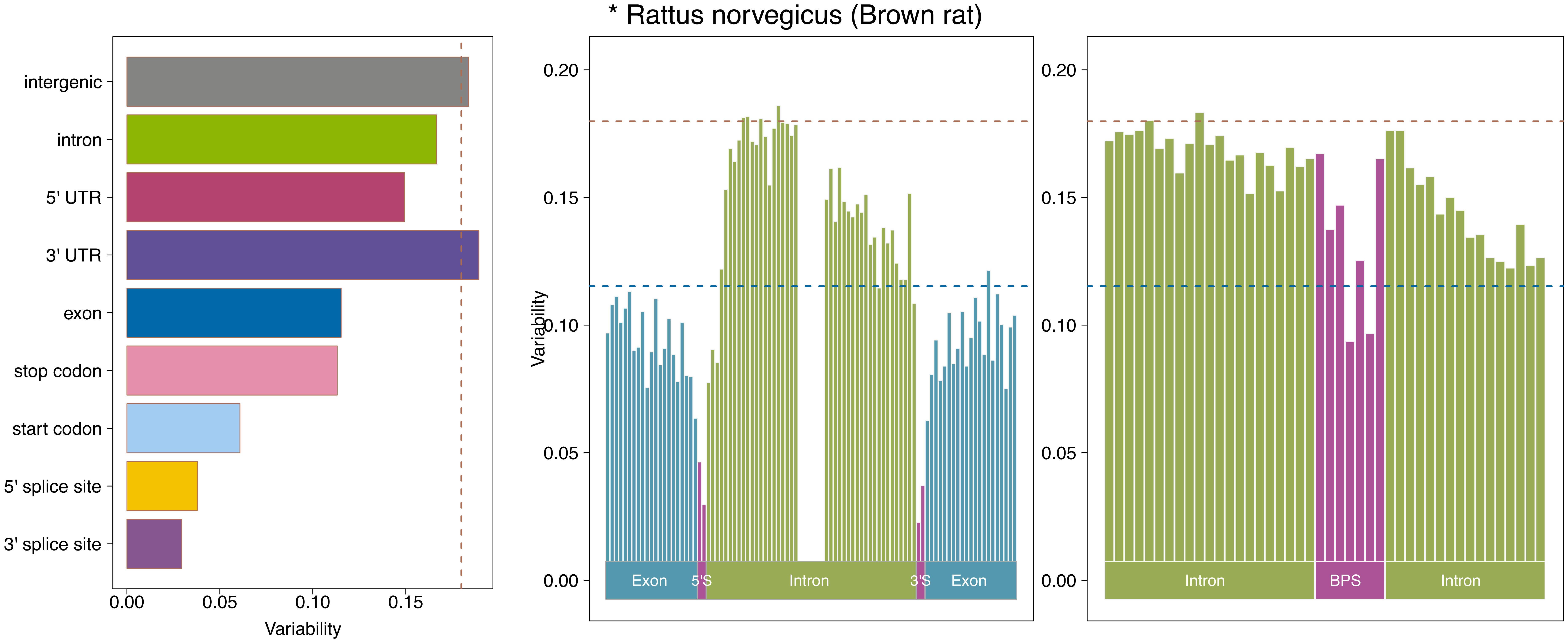
* Rattus norvegicus (Brown rat)
intergenic
intron
5' UTR
3' UTR
exon
stop codon
start codon
5' splice site
3' splice site
0.00
0.05
0.10
0.15
Variability
0.20
0.15
0.10
0.05
0.00
Variability
Exon
5'S
Intron
3'S
Exon
0.20
0.15
0.10
0.05
0.00
Intron
BPS
Intron

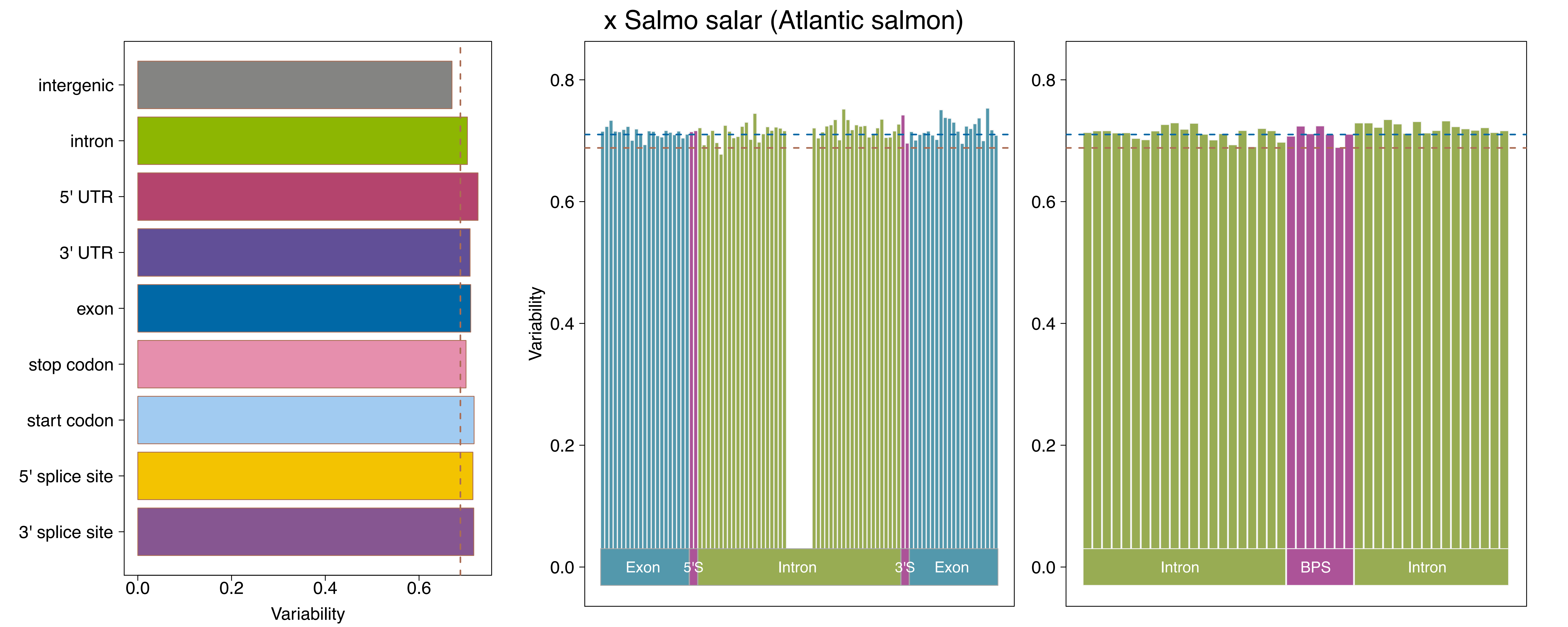

x Salmo salar (Atlantic salmon)
intergenic
intron
5' UTR
3' UTR
exon
stop codon
start codon
5' splice site
3' splice site
0.0
0.2
0.4
0.6
Variability
0.8
0.6
0.4
0.2
0.0
Variability
Exon
5'S
Intron
3'S
Exon
0.8
0.6
0.4
0.2
0.0
Intron
BPS
Intron

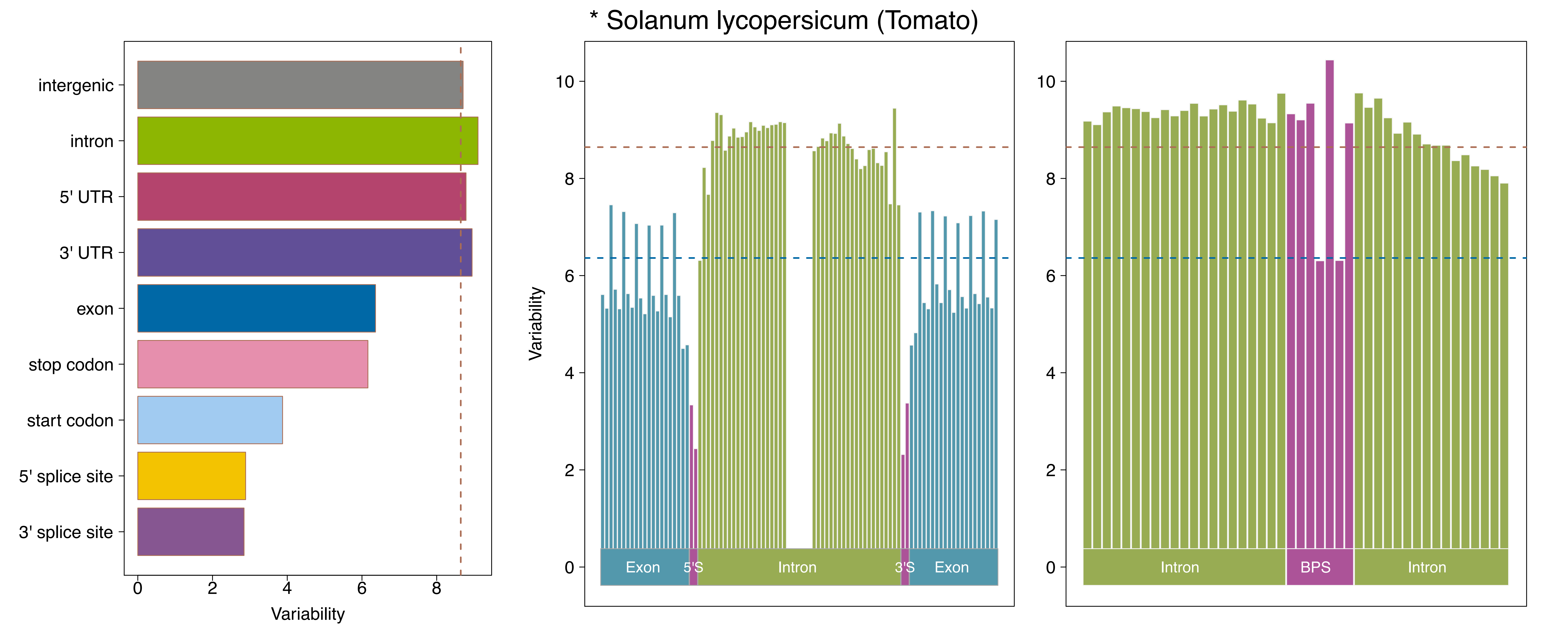

* Solanum lycopersicum (Tomato)
intergenic
intron
5' UTR
3' UTR
exon
stop codon
start codon
5' splice site
3' splice site
Variability
Exon
5'S
Intron
3'S
Exon
BPS
Intron

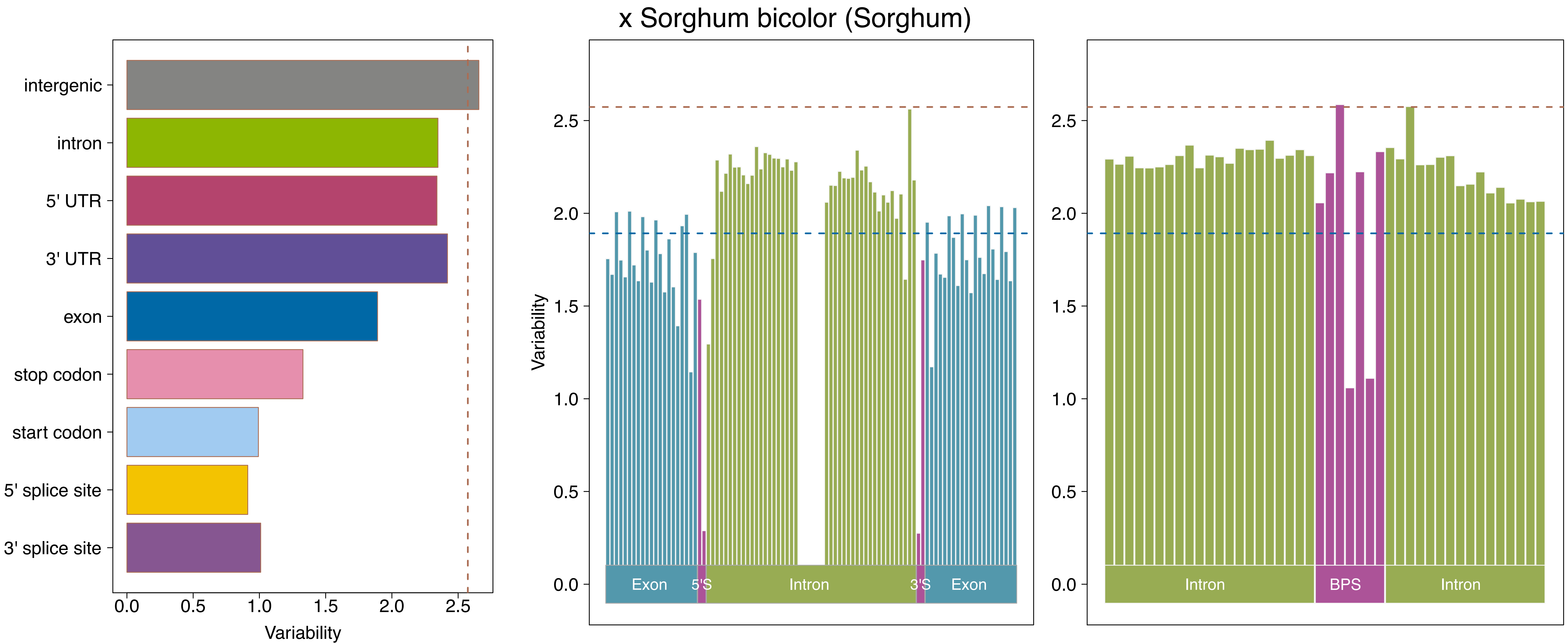
x Sorghum bicolor (Sorghum)
intergenic
intron
5' UTR
3' UTR
exon
stop codon
start codon
5' splice site
3' splice site
0.0
0.5
1.0
1.5
2.0
2.5
Variability
Exon
5'S
Intron
3'S
Exon
BPS
Intron

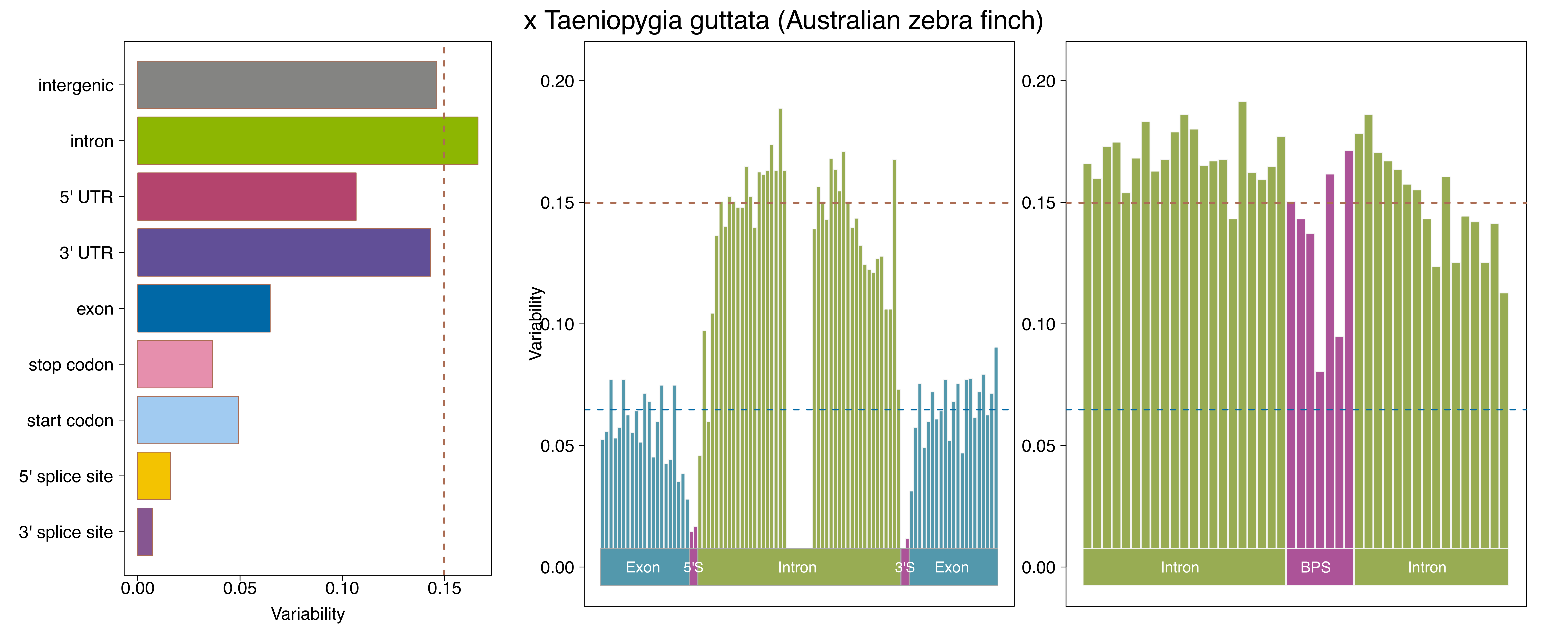

x Taeniopygia guttata (Australian zebra finch)
intergenic
intron
5' UTR
3' UTR
exon
stop codon
start codon
5' splice site
3' splice site
0.00
0.05
0.10
0.15
Variability
0.20
Variability
Exon
5'S
Intron
3'S
Exon
BPS

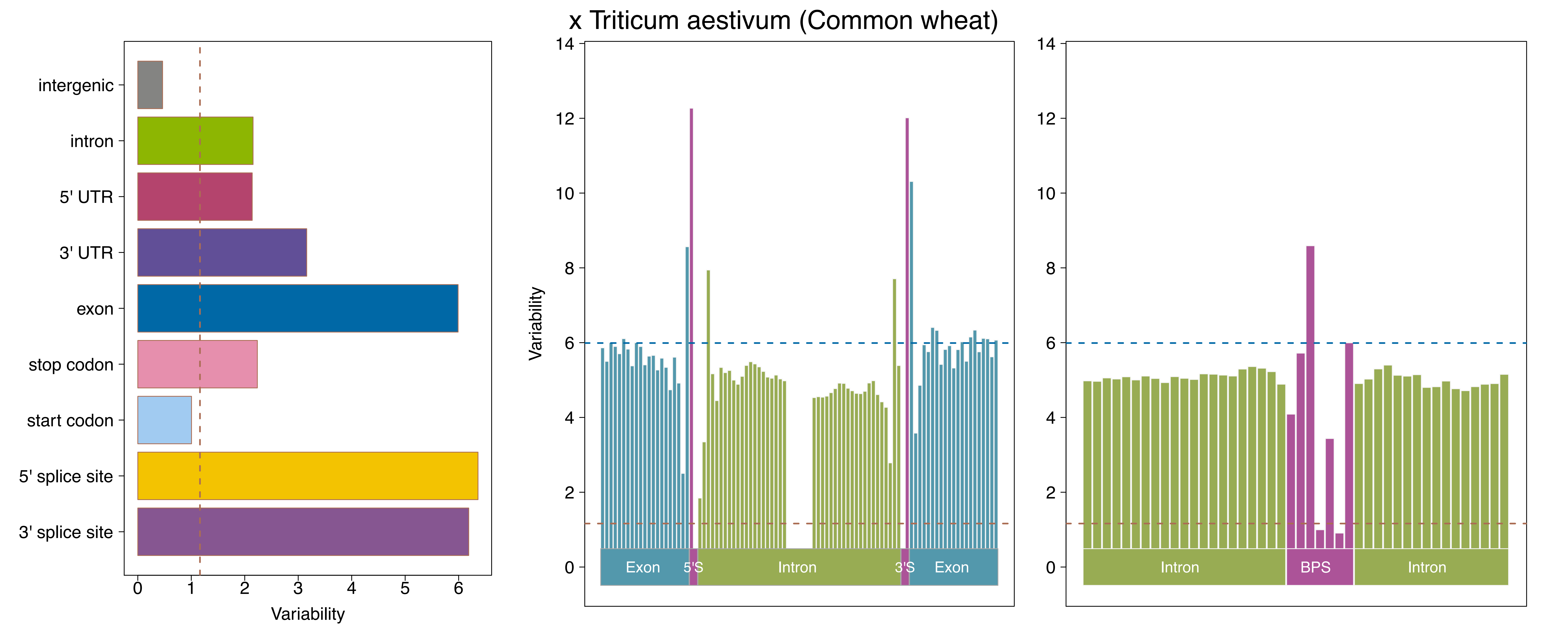
x Triticum aestivum (Common wheat)
intergenic
intron
5' UTR
3' UTR
exon
stop codon
start codon
5' splice site
3' splice site
0
1
2
3
4
5
6
Variability
Variability
14
12
10
8
6
4
2
0
Exon
5'S
Intron
3'S
Exon
Intron
BPS
Intron

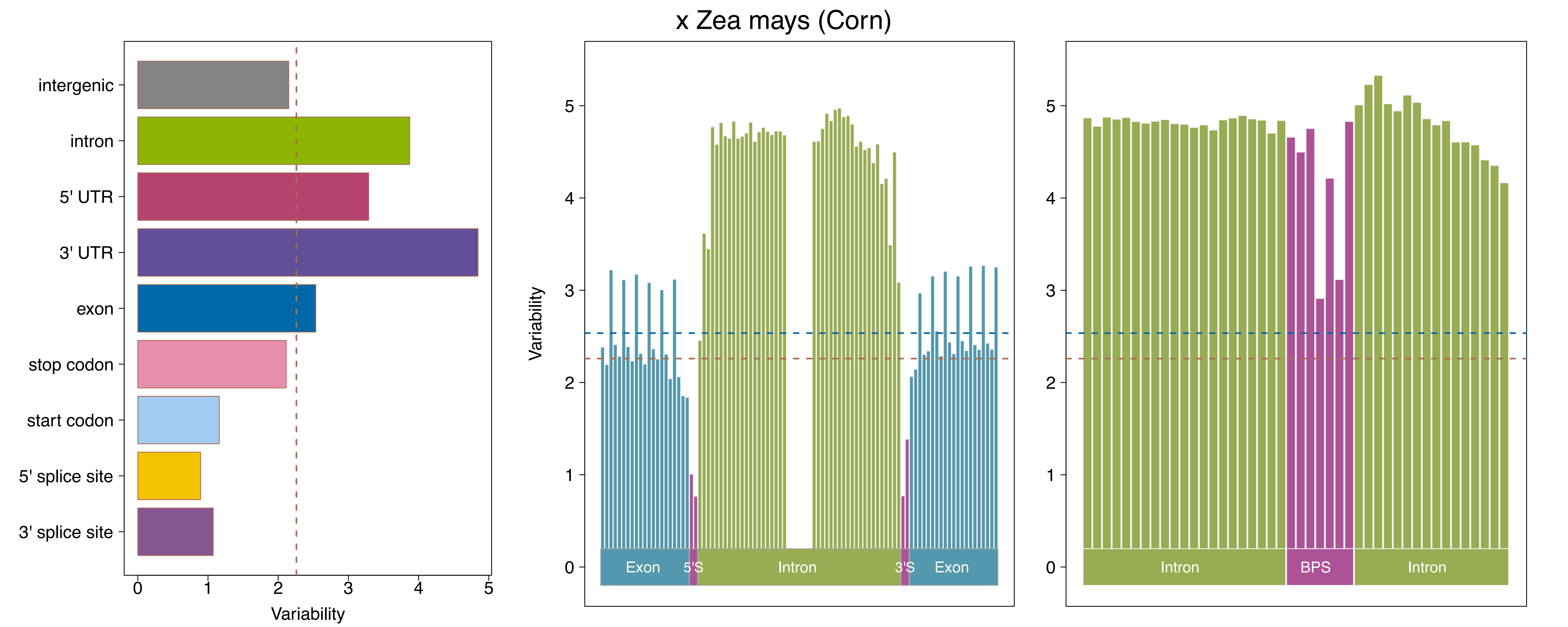

x Zea mays (Corn)
intergenic
intron
5' UTR
3' UTR
exon
stop codon
start codon
5' splice site
3' splice site
Variability
Variability
Exon
5'S
Intron
3'S
Exon
Intron
BPS
Intron
